# Inference and Validation of the Structure of Lotka-Volterra Models

**DOI:** 10.1101/2021.08.14.456346

**Authors:** Eberhard O. Voit, Jacob D. Davis, Daniel V. Olivença

## Abstract

For close to a century, Lotka-Volterra (LV) models have been used to investigate interactions among populations of different species. For a few species, these investigations are straightforward. However, with the arrival of large and complex microbiomes, unprecedently rich data have become available and await analysis. In particular, these data require us to ask which microbial populations of a mixed community affect other populations, whether these influences are activating or inhibiting and how the interactions change over time. Here we present two new inference strategies for interaction parameters that are based on a new algebraic LV inference (ALVI) method. One strategy uses different survivor profiles of communities grown under similar conditions, while the other pertains to time series data. In addition, we address the question of whether observation data are compliant with the LV structure or require a richer modeling format.

The code and data used in this manuscript are available at “https://github.com/LBSA-VoitLab/Inference_and_Validation_of_the_Structure_of_Lotka_Volterra_Models“.

## Introduction

Recent years have richly documented the enormous importance of microbiomes for human health and the well-being of the environment. In many cases, vast species diversity has been identified as a sign of normal operation, and substantial reductions in diversity have been associated with dysfunction or disease (*e.g*., [1, 2]). The compositions and specific roles of microbiomes are often difficult to comprehend, because the species within these communities tend to interact in complex ways, and the structure of the networks of interactions often changes over time. Addressing these challenges mandates computational approaches for inferring, characterizing, analyzing, and interpreting natural microbiomes and their responses to perturbations.

The default for characterizing a community of interacting microbial populations is a static network, which has the advantage of relatively straightforward graph-based analysis even if the number of species is large [3-8]. However, static networks have critical limitations, because their interactions are by default “symmetric” and time-invariant, which is often at odds with reality, where the effect of species A on species B is different than the effect of B on A, with the simple predator-prey system being an intuitive example [9, 10]. Also, the existence and strengths of interactions often change over time, as environmental systems, for instance, operate in seasonal cycles.

Addressing these issues requires dynamic mathematical representations, such as Lotka-Volterra (LV) systems [9-22] or discrete recursive Multi-Autoregressive (MAR) models [23-27]. Here, we focus on the former.

For *n* species, LV systems consist of *n* ordinary differential equations (ODEs) in the specific format

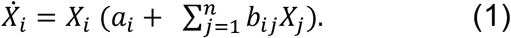

LV models have been employed to study macroscopic population interactions for almost one hundred years, and two-variable predator-prey models have traditionally been serving as examples and general motivation in introductory mathematical modeling classes for a long time. One reason is that it is straightforward to construct LV models from scratch, as the format for each population consists of one growth term (*a*_*i*_), one intraspecies interacting term (*b*_*ii*_) that is related to the “carrying capacity” *a*_*i*_*/b*_*ii*_, which limits the growth of this population as it becomes large, and one term each capturing a direct interaction of this population with possibly every other population in the system (*b*_*ij*_, *j≠i*). Of course, many of these direct interaction terms might be zero, but this information is typically not available at the beginning of a study.

One notes that these equations may be divided by *X*_*i*_, as long as these variables are not zero, which yields

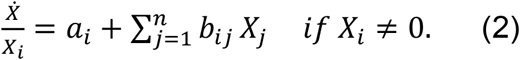

The left-hand side corresponds to the derivative of the logarithm of *X*_*i*_, demonstrating that changes in the logarithmic variables are defined by a linear system.

Although LV models have a long tradition of being considered appropriate default tools [28, 29] their format is sometimes criticized for being overly simplistic. This critique is often based on perception or bias, rather than proof or fact. In truth, simple models are often sufficiently accurate (although not 100% perfect), and they are typically much more robust than detailed mechanistic models, especially with respect to overfitting [30]. Examples of simple yet successful models in other contexts are growth laws, such as the Gompertz function that is heavily used in actuarial sciences [31], and statistical distributions, such as the normal, even though many actual data cannot be negative. In comparison, LV models are incomparably more flexible, and it was proved with mathematical rigor that they can have arbitrarily complicated dynamics, which is evident from the fact that any differentiable nonlinearities can be equivalently represented in this format if it includes auxiliary variables that are modeled in the same format [29, 32-34].

A different line of support comes from LV generalizations [35] in the format

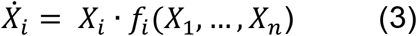

where the functions *f*_*i*_ are continuously differentiable. For many analyses of this format, one studies the (linearized) Jacobian of the system [35], which demonstrates that the traditional LV structure is, at least approximately, a reasonable representation for a much larger domain of functional forms.

This article describes four advances in the application of LV models to mixed community data. First, we introduce an algebraic LV inference (ALVI) method that can be used to infer interaction parameters from data on survivors of community experiments. Specifically, we address communities that are observed under similar environmental or experimental conditions but ultimately exhibit different survivor profiles. An example for the divergence is the observation that the same initial microbiome, under the exact same conditions—such as exposure to an antibiotic—may trend toward distinctly different end states; in other words, replicate experiments demonstrate that one or more different species are driven to extinction [36]. In fact, an LV model with ten species possesses over 1,000 (2^10^) possible steady states, which corresponds to the eventual coexistence of a subset of species that, in turn, directly corresponds to different survivor profiles. Another reason for differences in survivorship might stem from slightly different initial compositions of the community and, in particular, cases where one or more species are initially present with very low abundance—or even missing—within a set of experiments. In an experimental setting, it is furthermore possible to generate different outcomes by “spiking” a microbiome with an inoculum of some additional species, which subsequently might lead to different elimination scenarios [37-39]. Finally, one may create a series of survival experiments in which initially one or more microbial species are missing in order to discover new steady-state profiles.

In the second section of this paper, we slightly retool ALVI for a quantification of parameter values from time-series data that consist of species abundances throughout the history of a microbial community. The novel aspect of this estimation technique is that ALVI executes the inference with algebraic means.

The third advance described here is the use of ALVI as a diagnostic tool for assessing *a priori* to what degree the LV model structure adequately represents time series data. While the structure of any model is somewhat dictated by the data [40], it is quite rare that a modeler is able to confirm or refute the adequacy of a mathematical modeling format directly from observation data, but we show that such a validation assessment is indeed possible with ALVI.

Both the ALVI inferences from time series data and the LV-compliance analysis greatly benefit from the prior smoothing of the experimentally determined time course data. To this end, we introduce a strategy for determining effective settings for denoising raw time series data. We use for this purpose smoothing splines that subsequently replace the raw time series data and make ALVI faster and improve its results.

An important aspect of ALVI is its scalability. Most parameter estimation methods, such as gradient methods, scale rather poorly, even if many data are available, with some suffering severely from combinatorial explosion. By contrast, ALVI is based on decoupled systems that focus on one variable at a time and on methods of matrix algebra, which are known to scale very well in comparison to search algorithms.

## Methods

### 1. Algebraic parameter inference from survivor profiles

Most traditional estimation techniques for dynamical systems depend on time series measurements of population abundances within a community. If these are not available, it is presently difficult to assess the interaction structure of the community. The proposed algebraic LV inference (ALVI) offers a method for this situation. Specifically, it uses different steady-state species compositions of a community in response to natural variability or slightly different treatments. The concepts of this approach are as follows.

These interaction parameters *b*_*ij*_ (*i, j* = 1, …, *n*) of an LV model form a square matrix **B**. If the rank of **B** is full, the LV system has 2^*n*^ steady states. One of these corresponds to the only internal steady state, where all variables are non-zero, whereas all others contain at least one zero-valued variable. If the rank of the matrix of interaction parameters *b*_*ij*_ is not full, the steady-state profiles form a hyperspace or only allow an averaged solution obtained by regression.

In the context of bacterial microbiomes, the unique non-trivial steady state of total coexistence, if stable, may be interesting biologically, but it is less interesting mathematically because it attracts all trajectories that start from states without zeros, due to global stability. By contrast, a non-stable internal steady state has with a separatrix structure that subdivides the space into various basins of attraction.

The parameters *a*_*i*_ in Eq. (1) represent the intrinsic growth rates of the various species. For ease of discussion, we suppose that all *a*_*i*_ ≠ 0. Under opportune conditions, they may be inferred directly from independent monoculture experiments. Focusing exclusively on steady states, we may use these measured values or initially set all *a*_*i*_ = 1 and subsequently rescale all *b*_*ij*_ by the corresponding intrinsic growth rates, which yields the rescaled linear system

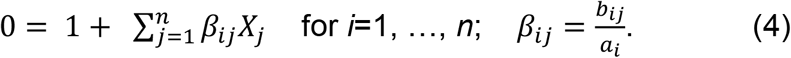

For ease of discussion, we consider a microbial community consisting of four species; much larger communities are addressed in exactly the same manner. Suppose for now that results are available from five survival experiments. We will later discuss cases with more or fewer experimental datasets. The differences in survivor profiles may be due to an unstable internal steady state and a separatix structure within the community that defines different basins of attraction. Alternately, different survivor profiles may be the consequence of including experiments where one or more species were missing from the beginning.

Among the five independent different steady states, one possibly, but not necessarily, consists of the only internal steady state, while in the other four cases at least one population is extinct. One could also have *m* (<5) profiles including one zero and 5-*m* profiles with two or more zeros. We call these observed steady-state *survivor vectors* (*S*_*i*1_, …, *S*_*i*5_)^*tr*^ and sort them such that *S*_*ii*_ = 0 for *i* = 1, …,4. In the fifth vector, all, some or none of the states are non-zero.

Consider the case of *X*_2_ = 0 at the steady state. To reflect a steady state of the dynamic LV model, the second equation in Eq. (4) is already satisfied, because the steady-state condition 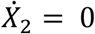 is achieved due to the fact that *X*_2_ = 0. However, the remaining three equations must be satisfied for each of the solution vectors. Formally:

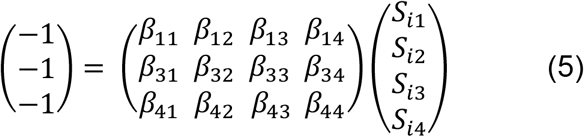

for *i* = 1, 3, 4, 5 and where *S*_*ii*_ = 0. Similar arguments hold if any of the other variables is zero. Thus, for each *i* in this example, we have three equations, yielding a total of twelve equations, and this number is equal to the number of unknown *β*_*ij*_.

The next task is expressing the interaction parameters *β*_*ij*_ as functions of the survivor vectors or, taken these together, of the survivor matrices. The task is accomplished by resorting all matrix equations as in (Eq. 5) for any *X*_*i*_ = 0. For instance, for *X*_1_ = 0 and *X*_2_ = 0, we obtain, respectively

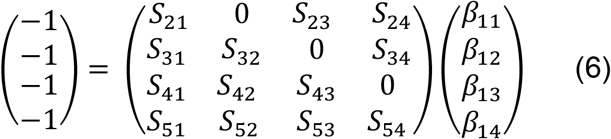

and

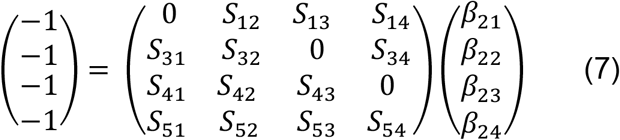

If the survivor matrices **S** have full rank, the solutions in terms of each vector of *β*_*ij*_ are obtained by simple matrix inversion. All *β*_*ij*_ parameter values are obtained in this manner, and rescaling retrieves the original *b*_*ij*_ in Eq. (1) if intrinsic growth rates are known. If they are not known, the interaction parameters are inferred except for one common scaling factor for each equation.

The types of solutions for the algebraic method depend on the number of survivor profiles. For a community of exactly *n* species and *n*+1 linearly independent profiles, matrix inversion yields a unique solution. If more than *n*+1 survivor vectors are available, two options are available. One may use some or all subsets of *n*+1 survivor profiles and use the collective results to form a model ensemble and study variability within this ensemble. Second, the overdetermined system may be solved in its entirety by means of a Moore-Penrose pseudo-inverse (left-inverse), which corresponds to the least-squares solution [41, 42]. Probably more pertinent is the situation of fewer datasets than variables are available, in which case the matrices, as in Eq. (6), are wide and cannot be inverted. One strategy is again the computation of a Moore-Penrose pseudo-inverse. The particular pseudo-inverse itself is not a good solution, because it minimizes the distance from the origin and therefore contains negative values. However, the total solution spans a space in which the optimal solution is located. *Supplement* Section 2 presents an example.

ALVI supposes that the survivor profiles constitute steady states, which has an important consequence. Namely, steady-state analyses by themselves do not allow assessments of the appropriate time scale. However, the most important inference from an LV model is usually the interaction structure within the community of populations, which is given by the interaction rates *b*_*ij*_ in Eq. (1) and *β*_*ij*_ in Eq. (4), which ALVI is designed to infer.

### 2. Algebraic parameter inference from time series data

Generic nonlinear parameter estimation methods for LV and other dynamic systems have been described in the literature many times and are therefore not discussed here (*e.g*., see reviews [43-45]). Instead, ALVI offers a novel, genuinely different method of parameter values from time series that is purely algebraic.

#### 2.1. Mathematical approach

The starting point of ALVI is Eq. (1). In a preliminary step (*Methods* Section 4), we use smoothing to reduce noise in the data, while retaining important features in the data. The smoothing splines we use for this purpose not only interpolate the data and address noise, but also permit us to compute slopes at as many timepoints as we wish, because the splines are explicit functions. Introducing abbreviations for the estimated slopes and for products of observed population abundances as 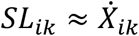 and *X*_*ijk*_ = *X*_*ik*_ · *X*_*jk*_ for species *i, j* = 1, …, *n* and timepoints *k* = 1, …, *K*, Eq. (1) becomes

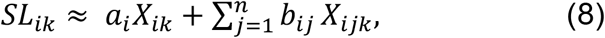

which is linear in the unknown parameters *a*_*i*_ and *b*_*ij*_ (*j* = 1, …, *n*). These approximate equations constitute a linear algebraic system of *n* blocks, where each block contains *K* equations, namely one per chosen timepoint.

The ALVI analysis may proceed in two ways. The first is straightforward linear regression with Eq. (8), which is further simplified by the fact that this regression can be executed for each block separately [46]. The second, novel option is matrix inversion. Specifically, for *n* dependent variables, we use *n*+1 readings of the dependent variables and the corresponding slopes to set up *n*^2^+*n* equations for calculating values of the *n*^2^+*n* unknown parameters. A simple example illustrates the approach.

Consider a hypothetical two-variable system, for which the population sizes and slopes in Table 1 had been obtained. Entering these values into Eq. (8) yields six equations in six unknowns:

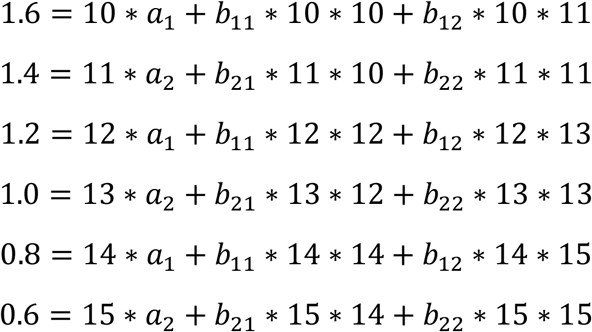

**Table 1:** Population sizes and slopes of a hypothetical dataset at different time points.

|  |  |  |
| --- | --- | --- |
| $t=1$ | $X_{11} = 10$ | $sl_{11} = 1.6$ |
| | $X_{21} = 11$ | $sl_{21} = 1.4$ |
| $t=2$ | $X_{12} = 12$ | $sl_{12} = 1.2$ |
| | $X_{22} = 13$ | $sl_{22} = 1.0$ |
| $t=3$ | $X_{13} = 14$ | $sl_{13} = 0.8$ |
| | $X_{23} = 15$ | $sl_{23} = 0.6$ |

Barring linear dependence, the system can be solved by straightforward matrix inversion.

A potential issue, which arises once in a while, is that this approach does not use the temporal structure of the data [47]. In other words, solving the linear system, ALVI provides estimates for the parameters of any LV system that reflects the observed values of variables and the corresponding slope values, independent of the specific timepoints where the system exhibits these values. For instance, in the above example, ALVI “does not know” that population sizes 10 and 11, along with slope values 1.6 and 1.4, were observed for *t*=1, and that population sizes 14 and 15 with slope values 0.8 and 0.6, were observed for *t*=3. As far as ALVI is concerned, these observations could have occurred in opposite order or at any other timepoints. Expressed in generic terms, ALVI does not account for the relative order of the sample points and the solution is therefore theoretically not unique. Also, if the chosen sample of points does not include the initial conditions, the estimated parameter set could therefore correspond to a system that does not pass through the observed initial points.

To counteract these issues, we introduce a preliminary step to choose appropriate sets of timepoints. This step is not mandatory and can be computationally intensive but finds the point sample that presents the best solution. First, we calculate a large number of combinations of timepoints to be used in ALVI. For an LV system with *n* dependent variables, we need (*n*+1) timepoints at which the *n* variables are observed. We could theoretically choose any values of variables and slopes from the smoothing splines, but as these have infinitely many points, this otherwise convenient feature here leads to a combinatorial explosion of choices. Instead, we use only those values of the smoothing function that have time coordinates coinciding with the raw data. This restricted choice usually leads to a reasonable number of combinations and can also serve as a reality check later. If there are not enough informative data points, the point set may be augmented with additional values from the splines. This situation could arise, *e.g*., if most of the data are steady-state measurements, where repeated observations are non-informative.

In the next step, an automated algorithm scans the chosen combinations of points, systematically feeding one point combination after another into ALVI. Using the parameter estimates from the ALVI solution with initial values from the smoothing function, we solve the Lotka-Volterra system numerically and calculate the SSE of the current estimate relative to the original data and to the smoothing function. We iterate these steps for every combination of points and at the end identify two parameter profiles that present the smallest SSEs, one pertaining to the original data and the other to the corresponding values of the smoothing function. These two sets are usually, but not always, the same.

For today’s typical data, it is unlikely that scanning through all combinations poses a computational challenge because inverting a matrix and solving the ODE system are both fast. Nonetheless, future microbiome techniques may generate much more comprehensive time-series data, in which case one might devise a customized some sampling scheme rather than addressing all combinations.

Two comments are worth adding: First, ALVI is based on matrix operations, which generally possess superb features of scalability, especially in contrast to gradient descent or evolutionary search algorithms, which typically slow down tremendously with increases in problem size. Obviously, ALVI is also immune to other issues of search algorithms, such as local minima. Second, this algebraic approach for time series directly permits inferences of growth and interaction parameters one equation at a time. This system decoupling amounts to an immense advantage for larger systems and adds to the favorable scalability of ALVI for time series data. It also suggests direct parallel execution.

### 2.2. Real-world data used for time series inferences

To illustrate ALVI for time series data, we analyze data from a recent study of four bacterial species [48], namely (with abbreviations we use) *Agrobacterium tumefaciens* (*At*), *Comamonas testosteroni* (*Ct*), *Microbacterium saperdae* (*Ms*), and *Ochrobactrum anthropi* (*Oa*). Biological details of the study are summarized in *Supplement* Section 3. As the specific dataset for our analysis, we use the community of all four species under the study’s control conditions. As Figure S8 of [48] indicates, all populations initially grow, but two populations begin to decline about six days into the experiments, presumably due to the depletion of nutrients.

We begin our analysis by smoothing the time course for each species and each replicate, as well as the average time course for each species. Because the absolute numbers of bacteria are quite large, we rescale the abundance data to units of 1,000,000 (see *Results* Section, Figs. 3 and 4).

## 3. A diagnostic tool for assessing the adequacy of the LV format

The ALVI approach in Section 1 offers an *a priori* validation method for assessing to what degree a given dataset is adequately captured by an LV model. The method is described and illustrated below for synthetic data that are LV-compliant and those that are not; an application to actual data is presented in the *Results* section.

To convey the overall concept, consider as a thought experiment the idealistic case that the data are noise-free and perfectly described by an LV system with *n* variables. Then, linearly independent blocks of values of the variables at any *n*+1 timepoints are sufficient to estimate all parameter values with perfect results. In fact, any such block will yield exactly the same parameter values. Turning the argument around renders it evident that the entire dataset is not well captured by a single LV model if different blocks of values at any *n*+1 timepoints yield substantially different parameter values. The method permits many variations in details.

Suppose a dataset consists of measurements over the time horizon *t* ∈ [*t*_*initial*_, *t*_*final*_]. It is again beneficial to subject the data to spline smoothing. Next, the portion of the dynamics is identified where the dependent variables have reached—or are very close to—the steady state. These data are removed, because sampling several points very close to the steady state is non-informative and tends to produce singular matrices, causing the system not to have a single, well-defined solution. Moreover, values close to the steady state pose a different challenge: due to their nature of polynomials, splines typically oscillate slightly near and at a steady state. These oscillations, even if small in magnitude, can introduce undue variations in the parameter estimates that mask the true signal distinguishing LV-compliant from non-LV-compliant datasets. It is therefore advisable to remove consecutive points of essentially the same value. A reasonable criterion might be the exclusion of values of the variables if they exhibit a slope and second derivative lower than a preset threshold, such as 10^−3^ in absolute value. There is no problem if some scattered points in the sample have low derivatives, for instance, at the peak of an overshoot, but points with low derivatives should not be consecutive with the steady-state block of points, which is typically reached at the end of the dataset. The result of this trimming step is often a reduced time horizon of *t* ∈ [*t*_*initial*_, *t*_*d*_], *t*_*d*_ < *t*_*final*_, where *t*_*d*_ is the last timepoint preceding the steady-state block at the end of the dataset.

Depending on the density of measurements, one chooses several intervals, which may or may not be overlapping. To avoid linear dependencies in the ALVI matrix, it is ill-advised to choose *n*+1 consecutive points for the subsample, if these are dense in time. Instead, after starting a subsample at point *k*, one might use every fifth, tenth or so consecutive point, such as *k, k*+5, *k*+10, *k*+15, …. Obviously, the details should be tuned to the data. Barring numerical exceptions due to unsuitable point choices, this subset of data immediately generates parameter inferences per matrix inversion. Now we slide a window throughout the interval [*t*_*initial*_, *t*_*d*_], at each step selecting *n*+1 timepoints and computing parameter values. As an alternative to moving a window throughout the useable time range, one may choose independent subsets of points and compare the inference results.

In either case, all inferences will yield essentially the same parameter values if the data conform to the LV property. Slight variations, not necessarily normally distributed, are to be expected, due to small differences between the spline and the true model, and to numerical inaccuracies incurred during the matrix inversion. By contrast, if the data cannot be modeled with the same LV system in a satisfactory manner, the inferred parameter values must be expected to change substantially for different subsets of points. Plotted against the order in which the different parameters were estimated, the values may even exhibit trends that are clearly distinct from the small variations exhibited if the data that are LV-compliant. If so, the trends might be used to assess the “distance from the LV format.” If this distance is substantial, the LV format is inadequate and should be generalized for a more adequate data fit. This generalization may be imposed on the entire system or on only one or a few equations (see Discussion). At any rate, given the variation in parameter estimates for datasets not well modeled by a single LV model, it is reasonable to assume that a comparison of the degree of variability in the parameter estimates can serve as a convenient tool for distinguishing between LV-compliant and non-LV-compliant datasets.

As an illustration, consider a synthetic system composed of six state variables. The first two (*X*_1_ and *X*_2_) are governed by LV equations and unaffected by the rest of the system. The third and fourth (*X*_3_ and *X*_4_) are also defined in LV format but influenced by all other variables, including those not in LV format, while the fifth and sixth variables (*X*_5_ and *X*_6_) have equations in generalized mass action (power-law) or Michaelis-Menten format, respectively, and are influenced by variables *X*_1_ and *X*_2_. Details of the model are shown in the *Supplements*. Using this mixed set of equations, we created 100 data points and, for clarity of the illustration, did not add noise.

Figure 1 displays the data superimposed with model trajectories produced with the parameter estimates obtained from different windows in the LV-compliance test. As expected, *X*_1_ and *X*_2_ have trajectories that in almost all cases are very close to the data that had been generated with LV equations; slight deviations are due to numerical inaccuracies incurred by matrix inversions. By contrast, the other variables show substantial deviations in fits, which reflect the influence of the non-LV components.

**Figure 1:**
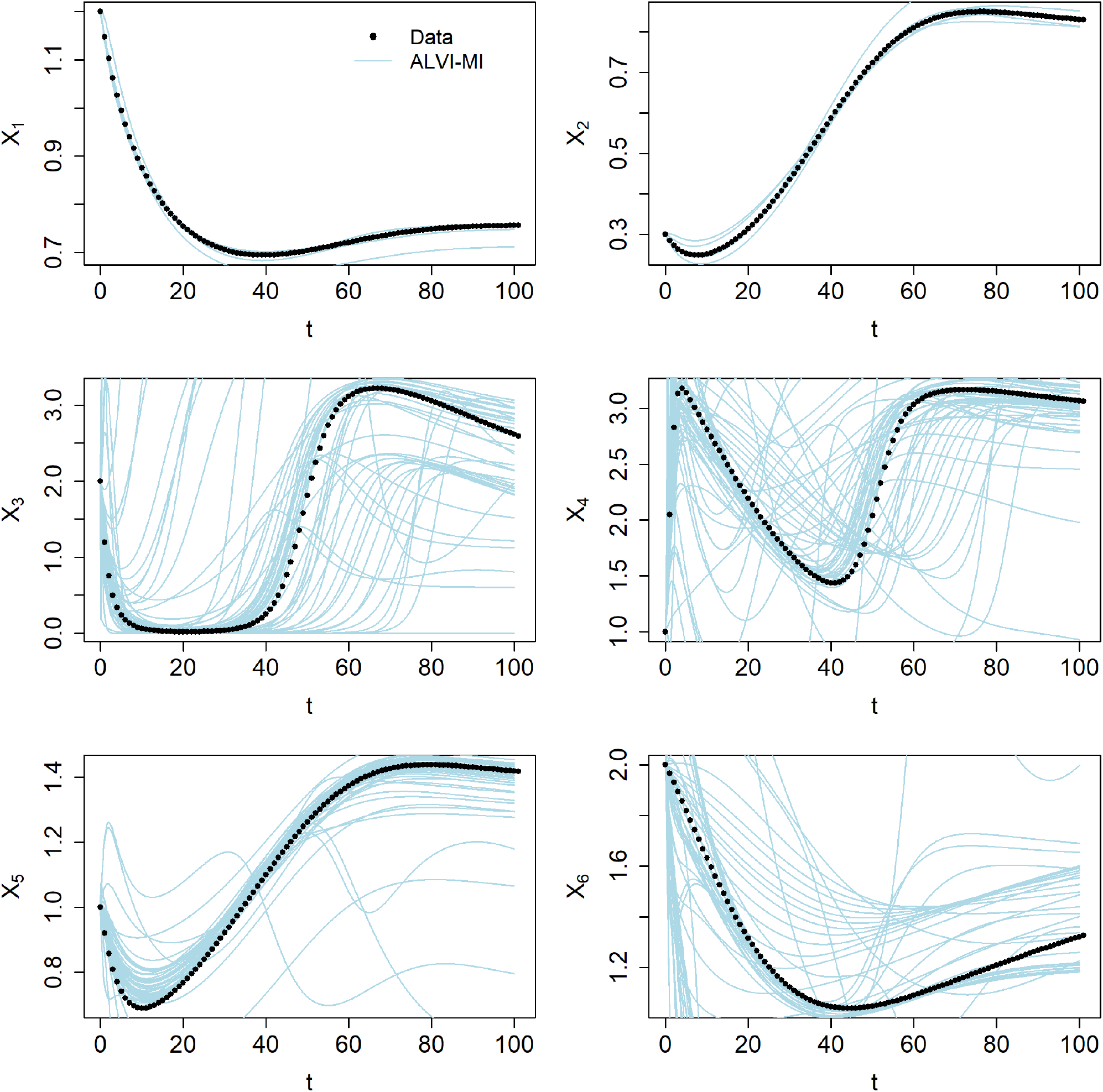
Results of an LV-compliance test with synthetic data. Data (dots) and trajectories (lines) were produced with sets of parameter estimates inferred for different positions of the sliding window in the LV-compliance test.

A semi-quantitative measure for LV-compliance is the analysis of variations in parameter values for different windows: If essentially the same LV model is inferred for every window, all variances are close to zero. In line with the visual results, the sets of estimates associated with each position of the sliding window are unchanging for *X*_1_ and *X*_2_, whereas they often exhibit substantial variations for *X*_3_ through *X*_6_ (Fig. 2).

**Figure 2:**
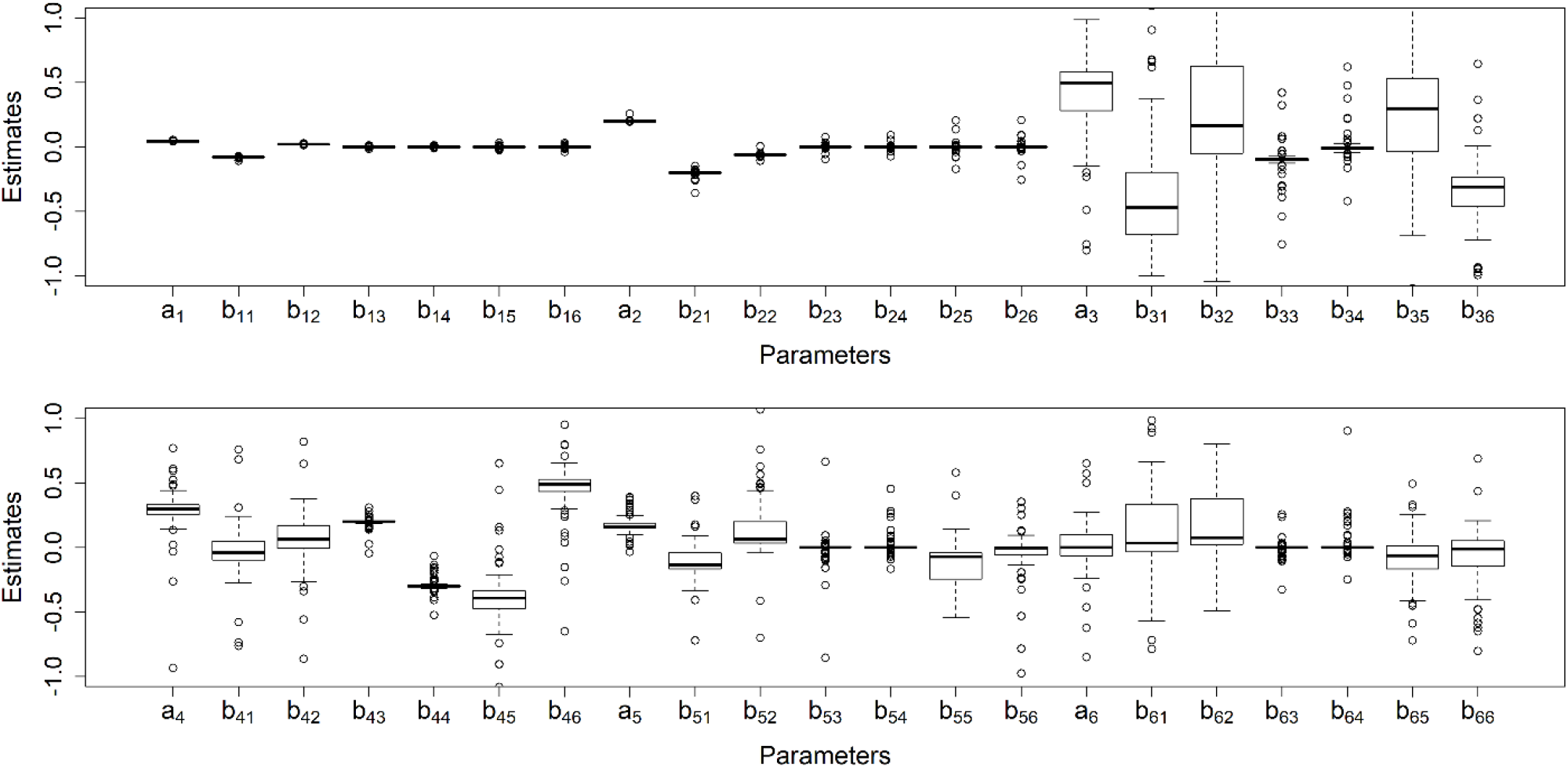
Variation in parameter values as indication of LV-compliance or non-compliance. Estimates and variations of parameters obtained in the LV-compliance test for different positions of the sliding window. The estimates associated with variables *X*_1_ and *X*_2_ display very little variation, in stark contrast to estimates associated with the other variables. For some parameters (*e.g*., *a*_3_ and *b*_31_), very large variations are obtained (see Fig. S2).

The result of the LV-compliance test is somewhat influenced by the degrees of freedom chosen for the smoothing splines. Given actual data, initial smoothing is always recommended to remove noise. For this illustration, we used 100 degrees of freedom, equal to the number of data points, although we typically recommend using the lowest degrees of freedom that return an adequate spline. This parsimony tends to prevent the emergence of undesirable artifacts.

## 4. Data smoothing

The earlier sections on algebraic parameter inferences from time series data require the estimation of slopes of the data trajectories at various timepoints. The latter task, in particular, is known to be strongly affected by noise. It is therefore useful to smooth the original raw data, while retaining important features in their time trends. This smoothing is not entirely trivial and unbiased, as the true nature of the data is seldom known and variations from a global trend line may be due to a true signal, noise, or a combination of the two. One could study the signal-to-noise-ratios throughout the experimental time period or apply outlier statistics in extreme cases, but a true distinction is impossible to state without bias, unless biological rationale suggests monotonicity or makes over- and undershoots unlikely, or if replicate data are available that exhibit over- and undershoot always at the same timepoints within the experimental time period.

Generically, data smoothing removes random variations in trends by decreasing the values of points that are higher than their closest neighbors and increasing them in the inverse situation [49]. Smoothing can be performed with a variety of methods [49-52]. Good general guidance for practical applications is provided in [49, 53]. Bin smoothers divide the data sample into bins and average the data in each bin. Moving average methods average the data in the form of a moving bin with a fixed number of data points instead of several static bins. If one applies a more complicated function to the moving bin, like a weighted moving average, one essentially uses a kernel density method. “Local regression” (LOESS) fundamentally is a weighted linear regression applied to the moving bin [51, 52].

We have had excellent success with splines [27]. In their original use, they are piecewise polynomial functions that pass through all sample points, are continuous, and have first and second derivatives, which are continuous at junction points between adjacent intervals. Here we are interested in smoothing splines, for which the first condition is substituted by a least-squares fit and balanced with an additional criterion that penalizes splines with high second derivative values that are indicative of local roughness [51, 52, 54]. The splines do not only address noise, they also permit the computation of slopes and the interpolation of the trends in dependent variables at as many timepoints as is desired.

Here, we mainly use splines, but alternative smoothing techniques could be employed. The only requirement is an effective estimation of the smoothed data points for the dependent variables, as well as their slopes. This output is then passed to the ALVI parameter estimator.

The first step for the use of a smoothing spline is the choice of its degrees of freedom, which dictate the level of smoothness of the spline. A natural default is a function that passes through every data point. Typically, this function is rather rugged but has a sum-of-squared-errors (SSE) value of 0. It is necessary to balance this perfect, rugged fit with a penalty for high values of second derivatives along each time course. It is possible to automate this balancing task, by calculating the trace of the smoothing matrix [55], a strategy that was automated, for instance, in the R package *mgcv*. However, idiosyncrasies of the datasets usually yield results that are not as satisfactory as those chosen by a knowledgeable human who is familiar with the characteristics of the data and has rationale to discern potential signal in the data from noise or observation error. Here, we determined a suitable degree of freedom manually, using as quality criterion a pleasing representation of the data.

If we opt to use LOESS, we are able to control the roughness in the smoothed time trends of the different variables. The R package *fANCOVA*, for example, facilitates this task, as it allows optimal control of the smoothing process. The user chooses the degree of the smoothing polynomial and selects an automatic smoothing parameter based on either a bias-corrected Akaike information criterion (AICC) or generalized cross-validation (GCV).

Once suitable smoothing is achieved that minimizes noise without compromising the true signal, the resulting smoothing function allows interpolation and the calculation of derivatives, which are passed to the ALVI parameter estimator.

## Results

### 1. Algebraic parameter inference from survivor profiles

For an illustration of the survivor-based variant of ALVI, suppose the system under investigation consists of four populations of different species. Thus, our starting point is the model equation

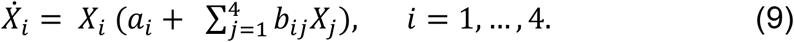

Suppose the matrix **B** of *b*_*ij*_’s of the system has full rank. The system then has a total of 16 steady states. One of these states represents full coexistence, where the value of each dependent variable is strictly positive, whereas at least one variable has a value of 0 in the remaining 15 steady states.

Because this method focuses on interaction parameters and because steady-state analyses do not allow assessments of the time scale of a system, we define the growth parameters *a*_*i*_ as 1 and rescale the results later, if time-scale information becomes available. For simplicity of notation, we rename *β*_*ij*_ = *b*_*ij*_/*a*_*i*_ for all *i* and *j* and obtain the steady-state equation

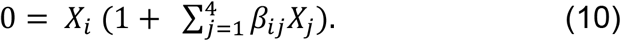

For a given *i*, Eq. (10) can be satisfied for two reasons: Either *X*_*i*_ is zero or the term in parentheses is zero. Importantly, if *X*_*i*_ =0, the term in parentheses does not have to equal zero for the system to be in a steady state.

We begin with the ideal case that the experiments had generated five independent survivor profiles; the method for fewer profiles is discussed in *Supplement* Section 2.

Suppose that the survivor vectors *S*_*k*_, *k* = 1, …, 5 represent independent, different steady states. As a numerical illustration, consider

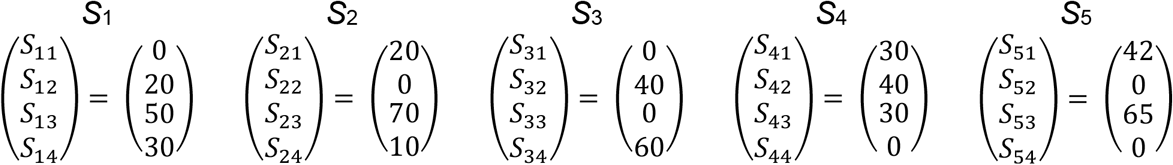

To begin the demonstration, we focus on *X*_1_ and use survivor vectors *S*_2_ to *S*_5_ to infer the interaction parameters *β*_11_, …, *β*_14_, which represent the effects of the populations 1, 2, 3, and 4 on population 1, respectively. Rearranging the equations in Eq. (10), as discussed in the *Methods* Section, we obtain

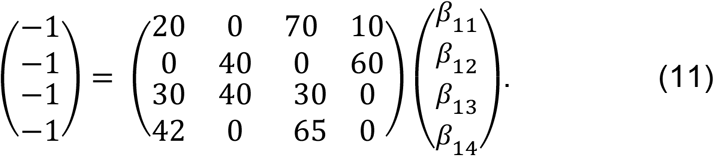

Inversion of the matrix and left-multiplication to the left-hand and right-hand sides directly yields

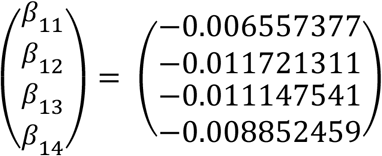

Similarly, we obtain the interaction parameters for populations 2, 3 and 4 as

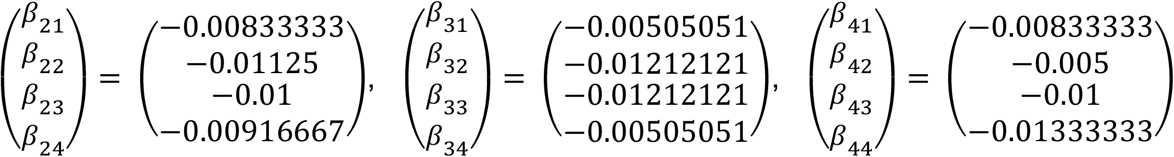

Entering these parameter values into an ODE solver demonstrates that all observed survivor profiles are steady states, with 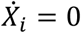 for all *i*.

### 2. Algebraic parameter inference from time series data

We apply the algebraic LV inference (ALVI) method to a community of four bacterial species, abbreviated as *At, Ct, Ms*, and *Oa* (for details, see the original article [48], *Methods* and *Supplement* Section 3). Three replicates are available for each population. To reduce numerical inaccuracies during the inference process, we scale all population sizes by 1/1,000,000. As a preliminary step, we submit each replicate for each species to denoising with smoothing splines (see *Methods*), and also compute splines for the means of the replicates. The computed splines are shown in Fig. S4. The degrees of freedom, along with subsamples for the replicates and means are presented in Table S2.

Using ALVI, we infer the intrinsic growth parameter for each species together with the interaction parameters from the smoothed time series data, using Eq. (1). The LV fits for all scenarios are shown in Fig. 3. Given the variability among the replicates, the data fits to the means can be considered adequate. The parameter values associated with each replicate and with their averages display consistency in signs in most, but not all cases (Table S3).

**Fig. 3:**
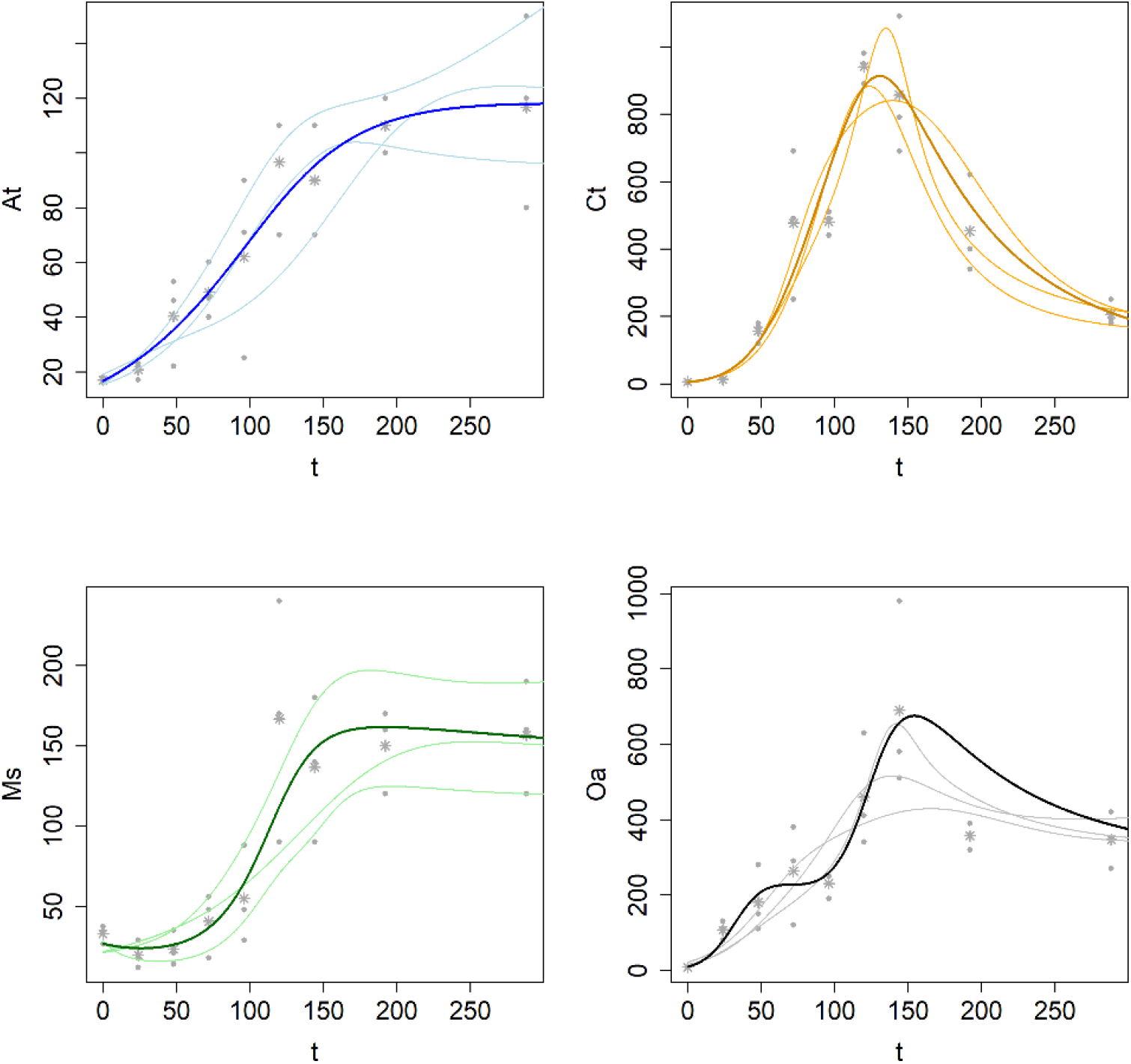
Data fits for all replicates and their means. The data were modeled with four-variable LV models, which were inferred from the splines in Fig. S4. Dots: raw data in units of 1 Million CFU/mL; 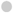: replicates; 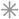: means of three replicates; light curves: fits of individual replicates; dark curves: fits of replicate means. Species abbreviations: *Agrobacterium tumefaciens* (*At*), *Comamonas testosteroni* (*Ct*), *Microbacterium saperdae* (*Ms*), and *Ochrobactrum anthropi* (*Oa*).

### 3. Diagnostics of the appropriateness of the LV format

A difficult question for every modeling effort is whether the mathematical format of the chosen model is adequate for the data of interest. Here we analyze this question by re-analyzing the bacterial community data from *Results* *Section* 2, using the averages of three replicates. The argument is as follows: If the LV format is representative for these data, then fitting any parts of the dataset should result in dynamic trends, as well as growth and interaction parameters that are very similar to those in fits of all data or of data in any other parts of the dataset.

As a coarse preliminary assessment of LV-compliance, we compare the quality of fit (SSE) for the first half of the data (*t* ∈ [0, 144]) with the fit to all data (*t* ∈ [0, 288]), in both cases using the means of replicates, but computed only for the data points from first half of the experiment (*t* ∈ [0, 144]). As before, we compute splines (results not shown) and then infer LV models from these splines. The results are shown in Fig. 4 and in Table 2. The fits for population *At* are almost indistinguishable, while the other three are somewhat different. Table 2 indicates that the SSE of the fit obtained for the entire experiment, but evaluated only for *t* ∈ [0, 144], is larger for replicates 1 and 3 as for the fit computed only for *t* ∈ [0, 144]. For the means and replicate 2 the SSEs are similar. In order to judge the significance of the difference between the two results, we compare the SSEs of means with those of the individual replicates, which demonstrates that the SSEs of the two means are actually more similar than those for the individual replicates. Thus, the data are not entirely LV-compliant, but the LV format does capture the dynamics of the four species to a reasonable degree. The parameter estimates inferred for the two cases are presented in Table S4.

**Figure 4:**
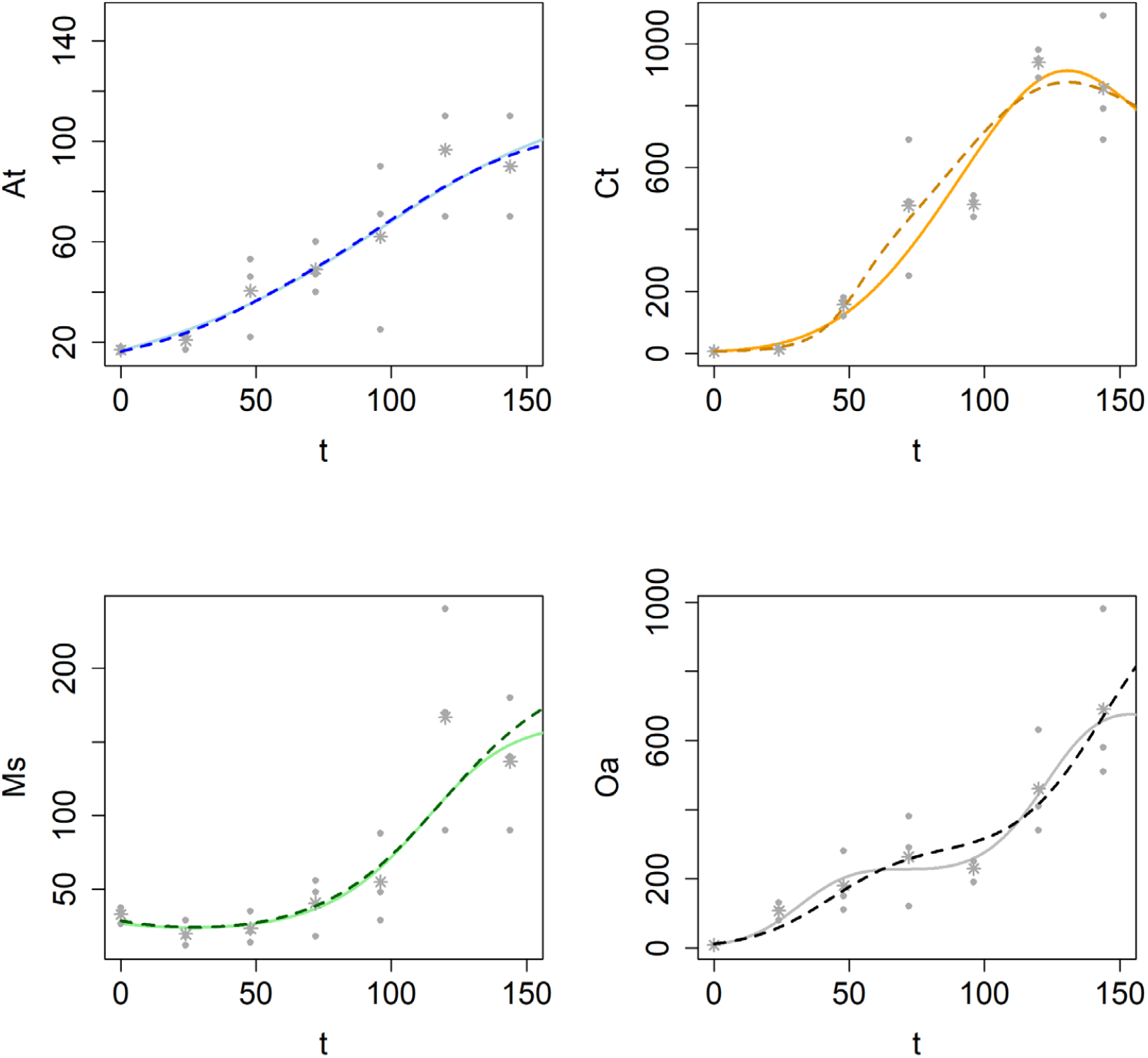
Alternative fits to bacterial community data. Means of the observed replicates of the dataset analyzed in *Results* Section 1 (Fig. 3) are fitted for the reduced time period of *t* ∈ [0, 144] (dashed lines) and compared to the earlier fits for *t* ∈ [0, 288] (solid lines). The fits are quite similar, although not identical. Dots: raw data in units of 1 Million CFU/mL;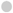: replicates; 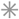: means of three replicates.

**Table 2:** Residual errors (SSEs) for data fits using all data or only the first half of the data. SSEs are presented for individual replicates and their means.

| | SSE of data fit for $t \in [0, 144]$ | SSE of data fit for $t \in [0, 288]$, but computed for time interval $[0, 144]$ |
| --- | --- | --- |
| Replicate 1 | 162,537 | 230,819 |
| Replicate 2 | 136,289 | 124,107 |
| Replicate 3 | 14,867 | 143,070 |
| Means of Replicates | 62,513 | 57,214 |

For a more formal LV-compliance test, we use ALVI with matrix inversion, as introduced in the *Methods* section. Because the data are only available at nine time points, we use all possible combinations of *n*+1 = 5 points up to timepoint *t*=144, rather than a sliding window, thereby creating an ensemble of models. If the data were truly LV-compliant, all samples would yield the same parameter estimates and data fits. Fig. 5 and Table S4 indicate that this is not the case, although the variation among models is much less than in the illustration of the *Methods* section. In particular, the trajectories for *At* and *Ms* are rather closely clustered, and differences could be due to numerical issues, as we found them in Fig. 1 for *X*_1_ and *X*_2_. There is less consistency for *Ct* and *Oa*, but by far not as much as for *X*_3_, …, *X*_6_ in Fig. 1. Overall, these findings reflect the result of the coarse preliminary analysis. Selecting the best solutions (with the lowest SSEs) among the sample of 21 fits creates a reasonable ensemble of models capturing the dynamics of the bacterial community.

**Figure 5:**
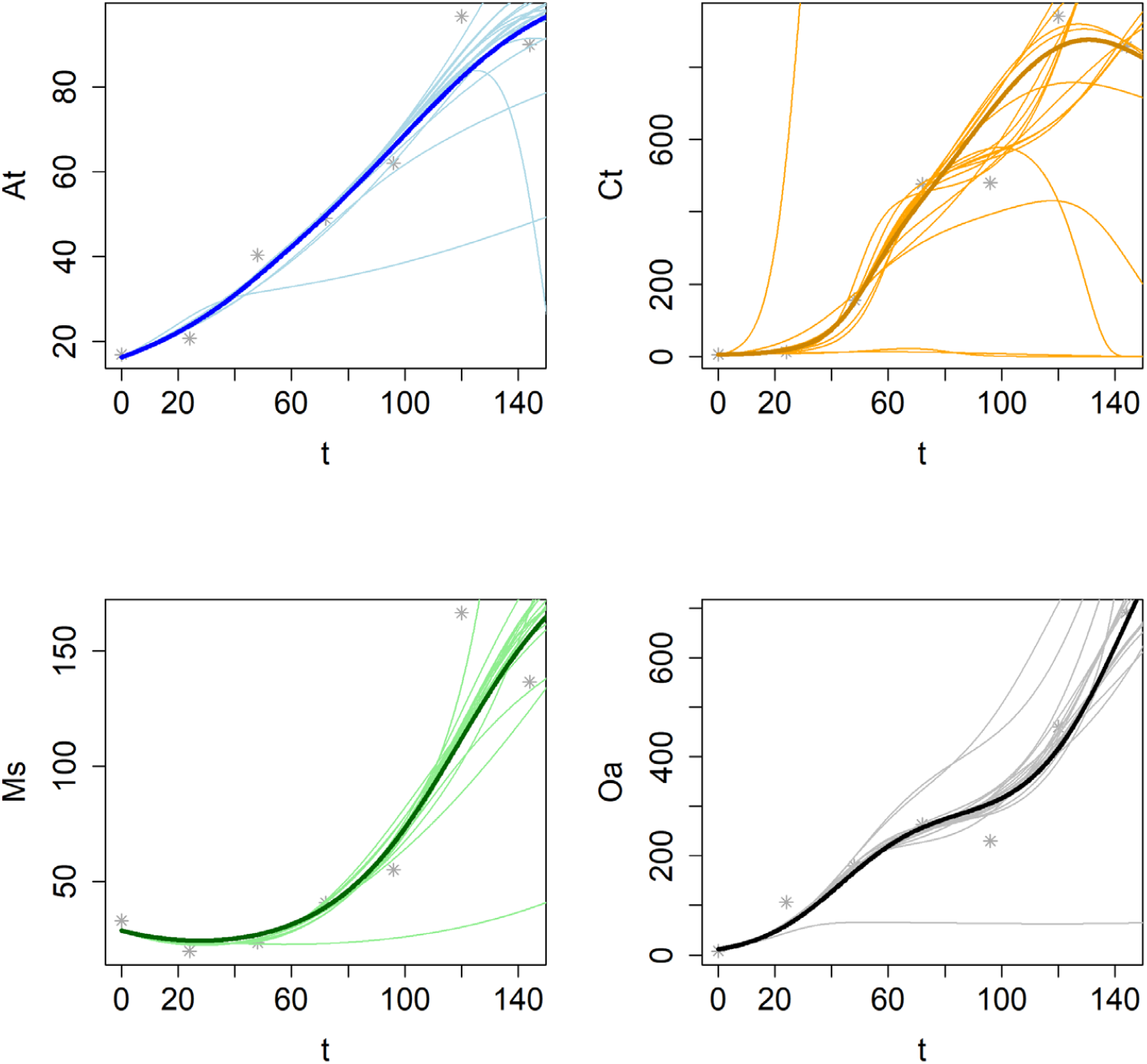
Ensemble of the 21 fits produced by parameter sets estimated from applying ALVI to all point sample combinations associated with timepoints 0, …, 144. Dots: raw data in units of 1 Million CFU/mL; 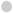: replicates; ✳: means of three replicates; light curves: fits with different parameter estimates; dark curves: best solution.

## Discussion

Lotka-Volterra models have been used for a long time. They are very easy to set up and interpret, and many model analyses are greatly facilitated by the fact that the equations, while nonlinear, have an almost linear character. In particular, the computation of steady states consists of simply solving a set of linear algebraic equations. In comparison to other nonlinear models, even the estimation of parameter values is relatively simple, if high-quality time series data are available. Nonetheless, tradition and expedience should not be the governing reasons to perpetuate the use of these models, and the nagging question remains: Are simple LV models rich enough to capture the complications of real-world population systems?

In a strict sense, the first answer must of course be ‘no,’ as LV models ignore many aspects of the complexity of life and are entirely deterministic. A better question, in line with George Box’s adage [56] that “all models are wrong but some are useful,” is whether LV models are useful, in particular for the new complex world of microbiomes, even if they are not 100% correct. In a generic sense, smaller, simpler models are often more robust to natural noise, variability, and perturbations, and they are most certainly less prone to overfitting [30]. At the same time, they may provide coarse-grained answers that might or might not be numerically optimal, but provide at least a qualitative or semi-quantitative impression of the interaction structure, which may actually be all that is of interest.

Here, we discuss two aspects of LV models. First, we address arguably the most important practical use of LV modeling, namely the inference of signs and magnitudes of interactions among coexisting populations. We demonstrate this aspect for two very different data types. Second, we propose a novel validation method for assessing to what degree the LV format is adequate or needs to be supplanted with a richer model structure.

The algebraic inference of LV parameters can be based on time series data or steady state survivor data and offers intriguing alternatives to existing methods. Concerning time series data, numerous methods have been proposed, including an efficient technique based on the estimation of slopes from data, which reduces the parameter inference to a straightforward method of linear regression [46]. The proposed ALVI method is rather similar in this respect, but permits inferences via matrix inversions, which are computationally very efficient and can naturally be used to create ensembles of well-fitting models if the data are at least approximately LV-compliant.

We have not encountered parameter inference approaches based on survivor profiles, even though somewhat similar thoughts were expressed in [35]. In addition to offering a novel approach to structure inference, the new availability of this very effective and readily scalable survivor-based method might trigger corresponding experimental designs for microbiome studies. For instance, it is often experimentally feasible *in vitro* to run a series of bacterial co-culture studies, where at first all species are included and then one species at a time is left out right from the beginning of the experiment. These experiments lead to a variety of survivor profiles, which form an excellent basis for the proposed ALVI method. It is at first surprising that interaction parameters can be inferred from steady-state data, and indeed ALVI generates results only up to a time-scaling factor. Specifically, the resulting parameters are in truth parameters scaled by the corresponding growth rates. Nonetheless, the most important information is gained anyway: because the growth rates are positive, the signs of interactions are retained, as are the relative magnitudes of the impacts of all species on the species in question.

For applications of ALVI to real data, it is highly recommended to use smoothing to reduce the effects of noise, before slope values are estimated from the data. Also, to be effective, ALVI requires data at more than *n*+1 timepoints, with *n* being the number of state variables. If so, an ensemble of models can be established by choosing different combinations of data points. Nonetheless, if fewer data are available, it is still possible to substitute or complement the set with points taken from the corresponding smoothing splines.

Because ALVI uses matrix inversion, the sample data points used to create the matrix must be linearly independent, which immediately indicates caution when using dependent variables close to a steady state.

The proposed diagnostic test of LV-compliance is a substantial and unusual advance, because it is rare that the appropriateness of a biomathematical model can be assessed without requiring numerous biological assumptions regarding the true mechanisms operating *in vivo*. Using an example from biochemistry, how can we objectively judge whether the ubiquitous Michaelis-Menten rate law is a “true” representation of an enzyme-catalyzed reaction *in vivo* [57, 58]? Similarly, an example from statistics is the frequent use of normal distributions even if the true variables are strictly positive.

To some degree, the interpretation of a LV-compliance test is a matter of judgment, as a decision must be felled regarding what is “close enough.” The results in Fig. 1 were clear-cut and consistent with the creation of the dataset used for the analysis. The decision for the real bacterial community data was less crisp.

The results of the LV-compliance test are affected by noise in the data, which permits choices in the level of smoothing. Although methods have been proposed to balance the goodness of fit with the smoothness of the time trends [55], as was pointed out in the *Methods* section, sound human judgement often achieves better results.

If the diagnostic test reveals that the data are indeed compatible with the LV format, the inferred parameter values are easily interpreted. Even in this case it is of course likely that other model formats could capture the same data with satisfactory accuracy [27]. Such alternative models can almost never be excluded. By contrast, if the diagnostic test reveals that the data are not compatible with the LV format, uncounted extensions are possible. A straightforward extension is the definition of environmental factors, whose effects on the system variables adhere to the format of LV equations. This type of generalization has been applied successfully to complex ecological systems [9, 10, 17].

The most natural generalization beyond this straightforward extension is the replacement of the simple multiplicative interaction terms with power-law terms containing either the two interacting species only, or these two species plus one or more additional modulating species. For example, the term *b*_*ij*_ *X*_*i*_ *X*_*j*_ in Eq. (1) could be replaced with the power-law term 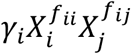, which allows greater flexibility modeling the particular effect of *X*_*j*_ on *X*_*i*_. An alternative is the term 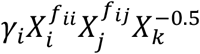, where the additional variable *X*_*k*_ has an inhibitory effect on the impact of *X*_*j*_ on *X*_*i*_, as is indicated by the negative power -0.5. In fact, it has been observed that a third species may alter the interaction structure among the two interacting species in a positive or negative fashion [36]. In this extended format, every variable of the system can potentially affect another variable simultaneously in multiple ways.

The generalization toward power-law functions directly leads to the Generalized Mass Action (GMA) format

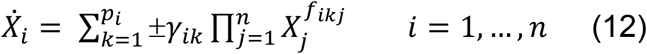

[59, 60], which has been used widely in biochemical and metabolic pathway studies and encompasses the LV format as a special case. In Eq. (12), *X*_*i*_ is affected by *p*_*i*_ interaction processes. Each contributing process has a rate constant (*γ*_*ik*_), which is multiplied to a product of variables directly influencing this process, and the strength of each influence is given by an exponent *f*_*ikj*_.

In cases where the diagnostic reveals that only some equations are not compatible with the LV format, the generalization toward GMA systems might be taken one term or one equation at a time, which retains the advantages of ALVI at least for compatible equations.

This GMA format is incomparably richer but loses many of the genuine advantages of the LV system. To some degree, the linearity of the process structure may be exploited for estimation purposes [61], but nothing comparable to the ALVI method is available.

## References

1. Delgado-Baquerizo, M., et al., Microbial diversity drives multifunctionality in terrestrial ecosystems. Nat Commun, 2016. 7: p. 10541.

2. Hahn, A., et al., Microbial diversity within the airway microbiome in chronic pediatric lung diseases. Infect Genet Evol, 2018. 63: p. 316–325.

3. Barberan, A., et al., Using network analysis to explore co-occurrence patterns in soil microbial communities. ISME J, 2012. 6(2): p. 343–51.

4. Chaffron, S., et al., A global network of coexisting microbes from environmental and whole-genome sequence data. Genome Research, 2010. 20(7): p. 947–959.

5. Faust, K., et al., Microbial Co-occurrence Relationships in the Human Microbiome. PLoS Computational Biology, 2012. 8(7).

6. Friedman, J., L.M. Higgins, and J. Gore, Community structure follows simple assembly rules in microbial microcosms. Nat. Ecol. Evol., 2017. 1: p. 0109.

7. Gilbert, J.A., et al., Defining seasonal marine microbial community dynamics. ISME J, 2012. 6(2): p. 298–308.

8. Ruan, Q.S., et al., Local similarity analysis reveals unique associations among marine bacterioplankton species and environmental factors. Bioinformatics, 2006. 22(20): p. 2532–2538.

9. Dam, P., et al., Dynamic models of the complex microbial metapopulation of lake mendota. Nature PJ Syst. Biol. Appl., 2016. 2: p. 16007.

10. Dam, P., et al., Model-based comparisons of the abundance dynamics of bacterial communities in two lakes. Sci. Rep., 2020. 10(1): p. 2423.

11. Lotka, A., Elements of Physical Biology. 1924 (reprinted as ‘Elements of Mathematical Biology’. Dover, New York, 1956): Williams and Wilkins, Baltimore.

12. Volterra, V., Variazioni e fluttuazioni del numero d’individui in specie animali conviventi. Mem. R. Accad. dei Lincei., 1926. 2: p. 31–113.

13. Fisher, C.K. and P. Mehta, Identifying keystone species in the human gut microbiome from metagenomic timeseries using sparse linear regression. PLoS One, 2014. 9(7): p. e102451.

14. Hanly, T.J., M. Urello, and M.A. Henson, Dynamic flux balance modeling of S. cerevisiae and E. coli co-cultures for efficient consumption of glucose/xylose mixtures. Appl Microbiol Biotechnol, 2012. 93(6): p. 2529–41.

15. Marino, S., et al., Mathematical modeling of primary succession of murine intestinal microbiota. Proc Natl Acad Sci U S A, 2014. 111(1): p. 439–44.

16. Mounier, J., et al., Microbial interactions within a cheese microbial community. Applied and Environmental Microbiology, 2008. 74(1): p. 172–181.

17. Stein, R.R., et al., Ecological modeling from time-series inference: insight into dynamics and stability of intestinal microbiota. PLoS Comput Biol, 2013. 9(12): p. e1003388.

18. Balagadde, F.K., et al., A synthetic Escherichia coli predator-prey ecosystem. Molecular Systems Biology, 2008. 4.

19. Berry, D. and S. Widder, Deciphering microbial interactions and detecting keystone species with co-occurrence networks. Frontiers in Microbiology, 2014. 5.

20. Bucci, V. and J.B. Xavier, Towards Predictive Models of the Human Gut Microbiome. Journal of Molecular Biology, 2014. 426(23): p. 3907–3916.

21. Fujikawa, H. and M.Z. Sakha, Prediction of competitive microbial growth in mixed culture at dynamic temperature patterns. Biocontrol Sci, 2014. 19(3): p. 121–7.

22. Liu, F., Y.Z. Guo, and Y.F. Li, Interactions of microorganisms during natural spoilage of pork at 5 degrees C. Journal of Food Engineering, 2006. 72(1): p. 24–29.

23. Certain, G., F. Barraquand, and A. Gårdmark, How do MAR(1) models cope with hidden nonlinearities in ecological dynamics?. Methods in Ecology and Evolution, 2018. 9(9): p. 1975–1995.

24. Holmes, E.E., E.J. Ward, and K. Wills, MARSS: multivariate autoregressive state-space models for analyzing time-series data. The R Journal, 2012. 4(1): p. 11.

25. Hytti, H., R. Takalo, and H. Ihalainen, Tutorial on multivariate autoregressive modelling. J. Clin. Monitor. Comp., 2006. 20(2): p. 101–108.

26. Ives, A.R., Predicting the response of populations to environmental change. Ecology, 1995. 76(3): p. 926–941.

27. Olivença, D.V., J.D. Davis, and E.O. Voit, Comparison between Lotka-Volterra and multivariate autoregressive models of ecological interaction systems. submitted, 2021.

28. May, R.M., Stability and Complexity in Model Ecosystems. 1973: Princeton University Press.

29. Peschel, M. and W. Mende, The Predator-Prey Model: Do we Live in a Volterra World? 1986, Berlin: Akademie-Verlag.

30. Lever, J., M. Krzywinski, and N. Altman, Model selection and overfitting. Nat Methods, 2016. 13: p. 03–704

31. Olshansky, S.J. and B.A. Carnes, Ever since Gompertz. Demography, 1997. 34(3): p. 1–15.

32. Peschel, M. and W. Mende, Riccati Darstellung nichtlinearer Systeme und elnlge meßtechnische Konsequenzen. Messen-Steuern-Regeln, 1983. 25: p. 67–69.

33. Peschel, M. and W. Mende, A unified modelling concept for nonlinear systems with Lotka-Volterra equations. Syst. Anal. Model Sjmul,, 1984. 1: p. 17–26.

34. Voit, E.O. and M.A. Savageau, Equivalence between S-systems and Volterra-systems. Mathem. Biosci., 1986. 78: p. 47–55.

35. Xiao, Y., et al., Mapping the ecological networks of microbial communities. Nat. Comm., 2017. 18: p. 2042.

36. Varga, J.J., et al., Antibiotics drive competitive release of rare pathogens in a chronic infection microbiome model. bioRXive, 2021.

37. Cira, N., M.T. Pearce, and S.R. Quake, Neutral and selective dynamics in a synthetic microbial community. PNAS U.S.A., 2018. 115(42): p. E9841–E9848.

38. Glassman, S.I., et al., Decomposition responses to climate depend on microbial community composition. Proc Natl Acad Sci U S A, 2018. 115(47): p. 11994–11999.

39. Livingston, G., et al., Competition–colonization dynamics in experimental bacterial metacommunities. Nat. Comm., 2012. 3: p. 1234.

40. Voit, E.O., Models-of-data and models-of-processes in the post-genomic era. Math Biosci, 2002. 180: p. 263–274.

41. Moore, E.H., On the reciprocal of the general algebraic matrix. Bull. Amer. Mathem. Soc., 1920. 26(9): p. 394–395.

42. Penrose, R., A generalized inverse for matrices. Proc. Cambridge Phil. Soc., 1955. 51: p. 406–413.

43. Chou, I.C. and E.O. Voit, Recent developments in parameter estimation and structure identification of biochemical and genomic systems. Mathematical Biosciences, 2009. 219(2): p. 57–83.

44. Gennemark, P. and D. Wedelin, Efficient algorithms for ordinary differential equation model identification of biological systems. IET Syst Biol, 2007. 1(2): p. 120–9.

45. Mendes, P. and D. Kell, Non-linear optimization of biochemical pathways: Applications to metabolic engineering and parameter estimation. Bioinformatics, 1998. 14(10): p. 869–883.

46. Voit, E.O. and I.-C. Chou, Parameter estimation in canonical biological systems models. Int. J. Syst. Synth. Biol., 2010. 1(1): p. 1–19.

47. Voit, E.O., A First Course in Systems Biology. 2nd ed. 2017, New York, NY: Garland Science.

48. Piccardi, P., B. Vessman, and S. Mitri, Toxicity drives facilitation between 4 bacterial species. PNAS U.S.A., 2019. 116(32): p. 15979–15984.

49. Batista, A.B.J. and P.P. S., An approach to outlier detection and smoothing applied to a trajectography radar data. J. Aerosp. Technol. Manag. [online], 2014. 6(3): p. 237–248.

50. Banks, D., Statistics Lectures. Duke University: http://www2.stat.duke.edu/∼banks/218-lectures.dir/dmlect2.pdf, 2009.

51. Cleveland, W.S., “Robust Locally Weighted Regression and Smoothing Scatterplots”, Journal of the American Statistical Association, Vol. 74,, Robust locally weighted regression and smoothing scatterplots. J. American Statistical Association, 1979. 74(368): p. 829–836.

52. Loader, C., 2012, Smoothing: Local Regression Techniques, in Handbook of Computational Statistics. 2012. p. 571–596.

53. Buja, A., T. Hastie, and i.R. Tibshiran, Linear smoothers and additive models. The Annals of Statistics, 1989. 17(2): p. 453–555.

54. Garcia, D., Robust smoothing of gridded data in one and higher dimensions with missing values. Computational Statistics & Data Analysis, 2010. 54(4): p. 1167–1178.

55. Wood, S.N., N. Pya, and B. Säfken, Smoothing parameter and model selection for general smooth models. J. Amer. Stat. Assoc., 2016. 111(516): p. 1548–1563.

56. Box, G.E.P., Robustness in the strategy of scientific model building, in Robustness in Statistics, R.L. Launer and G.N. Wilkinson, Editors. 1979, Academic Press: New York. p. 201–236.

57. Savageau, M.A., Enzyme kinetics in vitro and in vivo: Michaelis-Menten revisited, in Principles of Medical Biology, E.E. Bittar, Editor. 1995, JAI Press Inc.: Greenwich, CT. p. 93–146.

58. Savageau, M.A., Michaelis-Menten mechanism reconsidered: implications of fractal kinetics. J Theor Biol, 1995. 176(1): p. 115–24.

59. Savageau, M.A. and E.O. Voit, Power-law approach to modeling biological systems; I. Theory. J Ferment Technol, 1982. 60(3): p. 221–228.

60. Voit, E.O., Biochemical Systems Theory: A review. Int. Scholarly Res. Network (ISRN – Biomathematics), 2013. Article 897658: p. 1–53.

61. Dattner, I., H. Ship, and E.O. Voit, Separable nonlinear least-square parameter estimation for complex dynamic systems. Complexity, 2020: p. Article ID 6403641.

