## Supplemental Methods for "Inference and Validation of the Structure of Lotka-Volterra Models"

The supplements consist of three sections. The first contains additional details regarding the execution of an LV-compliance test. The second discusses inferences of parameters from survivor profiles, if the problem is underdetermined, while the third discusses further results from the application of the proposed ALVI methods to a bacterial community consisting of four species, as described in [1] and the Text.

### 1. Model of a Synthetic System Used for the LV-Compliance Test

#### *1.1. Equations and Parameters*

The synthetic system for testing LV-compliance is composed of six state variables. The first two ($X_1$ and $X_2$) are governed by regular LV equations and unaffected by the rest of the system. The third and fourth ($X_3$ and $X_4$) are also defined in regular LV format but influenced by all other variables, including those not in LV format, while the fifth and sixth variables ($X_5$ and $X_6$) have equations in generalized mass action (power-law) or Michaelis-Menten format, respectively, and are influenced by variables $X_1$ and $X_2$. The model has the format below, with parameter values and initial values presented in Table S1.

$$
\begin{aligned}
\dot{X}_1 &= X_1\,(a_1 + b_{11}X_1 + b_{12}X_2) \\
\dot{X}_2 &= X_2\,(a_2 + b_{21}X_1 + b_{22}X_2) \\
\dot{X}_3 &= X_3\,(a_3 + b_{31}X_1 + b_{32}X_2 + b_{33}X_3 + b_{34}X_4 + b_{35}X_5 + b_{36}X_6) \\
\dot{X}_4 &= X_4\,(a_4 + b_{41}X_1 + b_{42}X_2 + b_{43}X_3 + b_{44}X_4 + b_{45}X_5 + b_{46}X_6) \\
\dot{X}_5 &= c_{51}X_1^{f_{51}} + c_{52}X_2^{f_{52}} + c_{55}X_5^{f_{55}} \\
\dot{X}_6 &= \frac{V_{max2}\,X_2}{K_{M2} + X_2} - \frac{V_{max1}\,X_1}{K_{M1} + X_1}
\end{aligned}
\qquad \text{(S1)}
$$

***Table S1:***
***Parameter Values and Initial Conditions of the Model for Testing LV-Compliance (Eq. S1)***

| | | | | | | | | | | | |
|---|---|---|---|---|---|---|---|---|---|---|---|
| $a_1$ | 0.044 | $a_2$ | 0.2 | $a_3$ | 0.5 | $a_4$ | 0.3 | $c_{51}$ | 0.156 | $V_{max1}$ | 0.14 |
| $b_{11}$ | -0.08 | $b_{21}$ | -0.2 | $b_{31}$ | -0.5 | $b_{41}$ | -0.05 | $f_{51}$ | -0.5 | $K_{M1}$ | 0.1 |
| $b_{12}$ | 0.02 | $b_{22}$ | -0.06 | $b_{32}$ | 0.16 | $b_{42}$ | 0.05 | $c_{52}$ | 0.35 | $V_{max2}$ | 0.16 |
| | | | | $b_{33}$ | -0.1 | $b_{43}$ | 0.2 | $f_{52}$ | 0.6 | $K_{M2}$ | 0.2 |
| | | | | $b_{34}$ | -0.01 | $b_{44}$ | -0.3 | $c_{55}$ | -0.4 | | |
| | | | | $b_{35}$ | 0.3 | $b_{45}$ | -0.4 | $f_{55}$ | 0.9 | | |
| | | | | $b_{36}$ | -0.3 | $b_{46}$ | 0.5 | | | | |
| $X_{10}$ | 1.2 | $X_{20}$ | 0.3 | $X_{30}$ | 2 | $X_{40}$ | 1 | $X_{50}$ | 1 | $X_{60}$ | 2 |

### *1.2. Results*

A summary of trajectories and variations in parameter values for different locations of a sliding window of data points used is presented in Fig. S1.

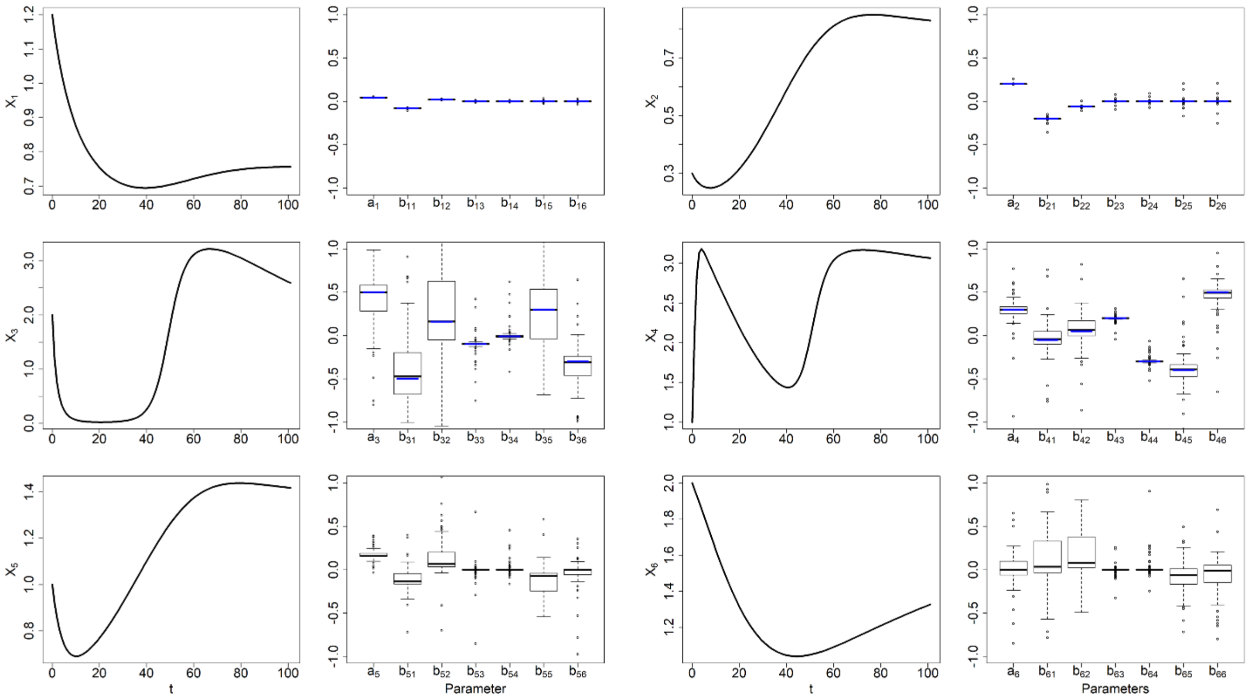


**Figure S1: LV-compliance test applied to a six-variable system (Eq. S1).** For each state variable, the model trajectory and variations in parameter values, as obtained from the LV-compliance test, are shown. The equations for $X_1$ and $X_2$ are in LV format and isolated form other variables; those for $X_3$ and $X_4$ are also in LV format but influenced by all variables of the system. The dynamics of $X_5$ and $X_6$ is modeled in generalized mass action (power-law) or Michaelis-Menten format, respectively; both are influenced by $X_1$ and $X_2$. Numerical details regarding this system are presented in Table S1. Strong variation in parameter estimates for different locations of the sliding window in the LV-compliance test indicates diminished adequacy of the LV format with respect to the particular data. Compliance is reflected in the narrow distributions of parameter values for $X_1$ and $X_2$, while the remaining distributions have much greater variance. Shown here are only estimates between -1 and 1. A more detailed representation of results is presented in Fig. S2.

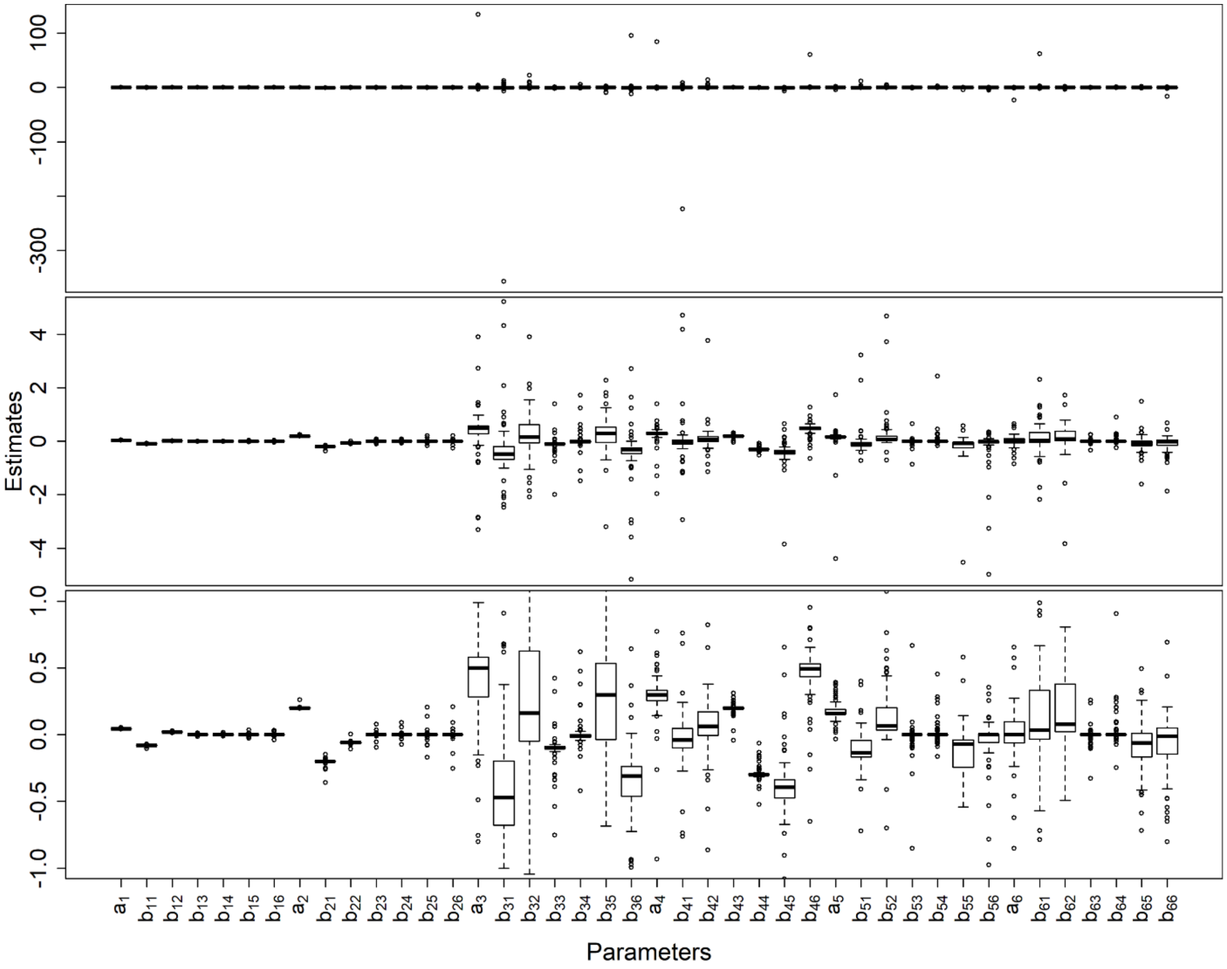


**Figure S2: Results of the LV-compliance test for the artificial model in Eq. (S1).** Each panel shows the variation in parameter values that result from computing an LV model for data in sliding a window (see Text and Fig. S1 for details). Small variation (parameter values associated with variables $X_1$ and $X_2$) indicates compliance with the LV format, whereas larger variations (for instance in $a_3$, $b_{31}$ and $b_{32}$) indicate deviations from this format. The three panels are associated with the same results, but have different vertical scales to offer a more comprehensive picture of parameter variability.

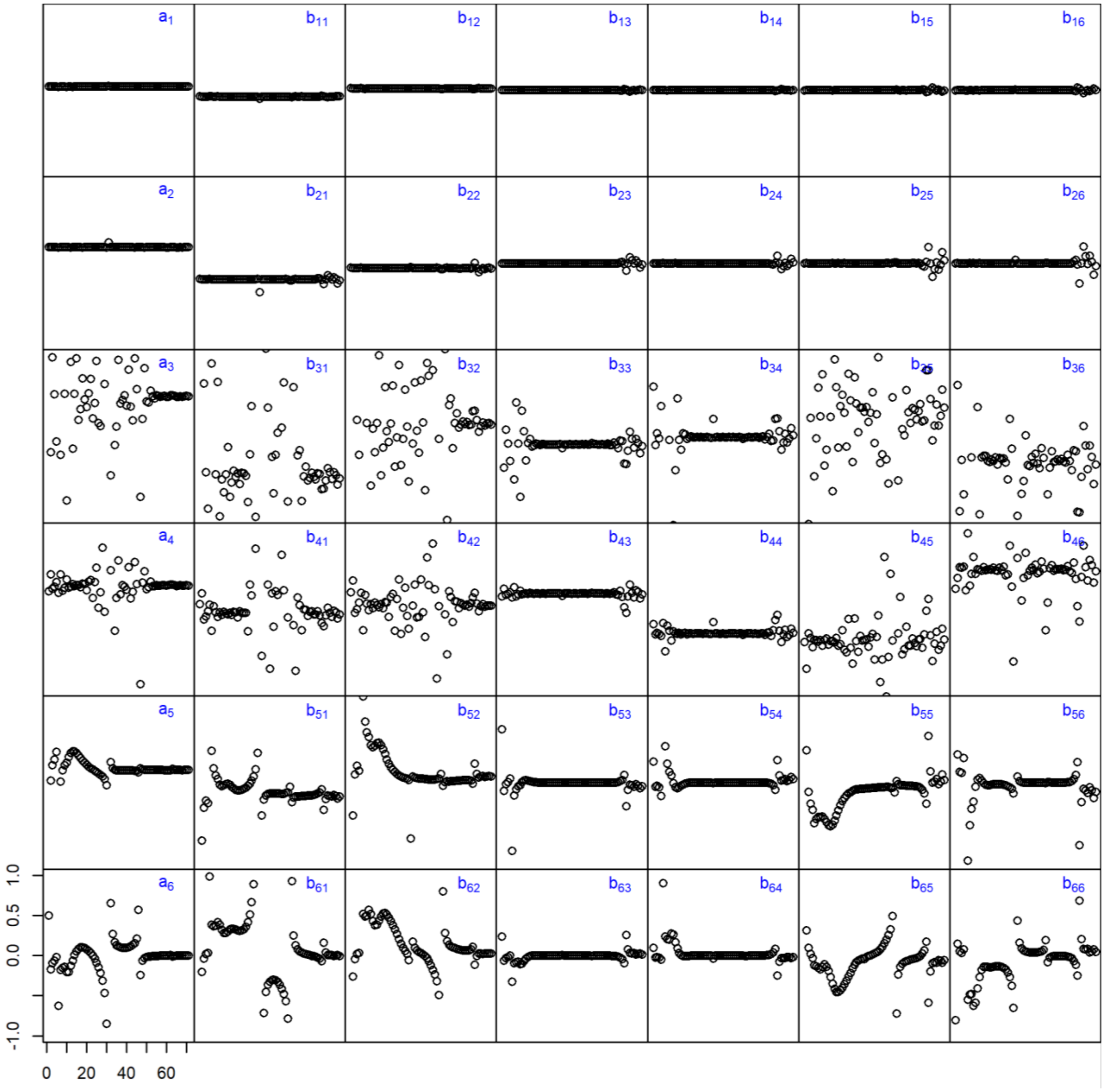


**Figure S3: Results of the LV-compliance test for the artificial model in Eq. (S1), displayed against the start location of the sliding window.** Trends in all parameters are shown for successive sliding-window samples. It becomes again clear that the equations for variables $X_1$ and $X_2$ (first two rows) are compliant with the LV format, whereas the other equations are not, even though some parameters exhibit small variations; an example is $b_{43}$. In order to provide a clear perspective of the different trends, only parameter estimates with values between -1 and 1 are shown.

### 2. Inference of Parameter Values from Fewer than *n*+1 Survivor Profiles

In the main Text, we discussed the case of exactly *n*+1 survivor profiles for a community of $n = 4$ species. Here, we analyze the same example, but suppose that the 5th survivor profile had not been observed, but that the other four profiles had the same abundances as before.

The equation for the reduced matrix $S_{R1}$, which equals the matrix in Eq. (11) in the Text without the last row, is

$$\begin{pmatrix}-1\\-1\\-1\end{pmatrix} = S_{R1}\begin{pmatrix}\beta_{11}\\\beta_{12}\\\beta_{13}\\\beta_{14}\end{pmatrix} = \begin{pmatrix}S_{21} & 0 & S_{23} & S_{24}\\S_{31} & S_{32} & 0 & S_{34}\\S_{41} & S_{42} & S_{43} & 0\end{pmatrix}\begin{pmatrix}\beta_{11}\\\beta_{12}\\\beta_{13}\\\beta_{14}\end{pmatrix} = \begin{pmatrix}20 & 0 & 70 & 10\\0 & 40 & 0 & 60\\30 & 40 & 30 & 0\end{pmatrix}\begin{pmatrix}\beta_{11}\\\beta_{12}\\\beta_{13}\\\beta_{14}\end{pmatrix}$$

The 4×3 matrix cannot be inverted, but its Moore-Penrose right pseudo-inverse is given as

$$S_{R1}^{+} = \begin{pmatrix}-0.001886792 & -0.003459119 & 0.011949686\\-0.009056604 & 0.003396226 & 0.017358491\\0.013962264 & -0.001069182 & -0.001761006\\0.006037736 & 0.014402516 & -0.011572327\end{pmatrix}.$$

Multiplied to the default growth-rate vector of 1s yields the set of *β*'s associated with $X_1$:

$$S_{R1}^{+}\begin{pmatrix}-1\\-1\\-1\end{pmatrix} = \begin{pmatrix}\beta_{11}\\\beta_{12}\\\beta_{13}\\\beta_{14}\end{pmatrix} = \begin{pmatrix}-0.006603774\\-0.011698113\\-0.011132075\\-0.008867925\end{pmatrix}.$$

The remaining *β*'s are computed in the same fashion:

$$\begin{pmatrix}\beta_{21}\\\beta_{22}\\\beta_{23}\\\beta_{24}\end{pmatrix} = \begin{pmatrix}-0.005862069\\-0.013034480\\-0.010000000\\-0.007931034\end{pmatrix}, \begin{pmatrix}\beta_{31}\\\beta_{32}\\\beta_{33}\\\beta_{34}\end{pmatrix} = \begin{pmatrix}-0.005172414\\-0.012068966\\-0.012068966\\-0.005172414\end{pmatrix}, \begin{pmatrix}\beta_{41}\\\beta_{42}\\\beta_{43}\\\beta_{44}\end{pmatrix} = \begin{pmatrix}-0.008571429\\-0.005714286\\-0.010000000\\-0.012857143\end{pmatrix}.$$

In this example, they are quite similar to the values obtained in the first section of the results in the main Text, but not exactly. In particular, the fifth profile, which was not used here, is almost but not exactly a steady state.

Recall that the $\beta_{ij}$ are interaction parameters, which are scaled by growth rates: $\beta_{ij} = b_{ij}/a_i$ for all *i* and *j*. If the growth rates $a_i$ are known, possibly from mono-culture experiments, rescaling yields estimates of the true interaction parameter values $b_{ij}$. But even if the $a_i$ are not known, the $\beta_{ij}$ still reflect the signs and the relative magnitudes of all interaction parameters associated with species *i*.

It is possible to attempt computing an internal steady state as $S_{int} = B^{-1}\,a$. However, this “survivor profile” contains negative values, namely:

$$\begin{pmatrix} S_{int1} \\ S_{int2} \\ S_{int3} \\ S_{int4} \end{pmatrix} = \begin{pmatrix} -666.6667 \\ -53.3333 \\ 133.3333 \\ 286.6667 \end{pmatrix}.$$

This computational result is biologically irrelevant, due to the negative values.

The above pseudo-inverse solutions are obtained from underdetermined systems and represent particular solutions within entire spaces of solutions. The general pseudo-inverse solution, here exemplified for population 4, is given by equations like

$$\begin{pmatrix} \beta_{41} \\ \beta_{42} \\ \beta_{43} \\ \beta_{44} \end{pmatrix} = S_{R4}^{+} \begin{pmatrix} -1 \\ -1 \\ -1 \end{pmatrix} + [I - S_{R4}^{+}\, S_{R4}]\, w,$$

where *I* is the identity matrix and *w* is an arbitrary vector. For our numerical example, we obtain

$$S_{R4}^{+} \begin{pmatrix} -1 \\ -1 \\ -1 \end{pmatrix} = \begin{pmatrix} -0.008571429 \\ -0.05714286 \\ -0.01 \\ -0.012857143 \end{pmatrix},$$

$$S_{R4}^{+}\, S_{R4} = \begin{pmatrix} -0.065 & 0.046428571 & 0.027142857 \\ 0.015 & -0.010714286 & 0.001428571 \\ 0.02 & 0 & -0.01 \\ -0.01 & 0.007142857 & 0.015714286 \end{pmatrix} \begin{pmatrix} 0 & 20 & 50 & 30 \\ 20 & 0 & 70 & 10 \\ 0 & 40 & 0 & 60 \end{pmatrix},$$

and

$$I - S_{R4}^{+}\, S_{R4} = \begin{pmatrix} 0.071428571 & 0.214285714 & 0 & -0.142857143 \\ 0.214285714 & 0.642857143 & 0 & -0.428571429 \\ 0 & 0 & 0 & 0 \\ -0.142857143 & -0.428571429 & 0 & 0.285714286 \end{pmatrix}.$$

For instance, the choice

$$w = \begin{pmatrix} 1 \\ 0 \\ 0 \\ 0 \end{pmatrix}$$

determines the 4$^{th}$ row vector of a as

$$\begin{pmatrix}\beta_{41}\\ \beta_{42}\\ \beta_{43}\\ \beta_{44}\end{pmatrix} = \begin{pmatrix}-0.008571429\\ -0.05714286\\ -0.01\\ -0.012857143\end{pmatrix} + \begin{pmatrix}0.071428571 & 0.214285714 & 0 & -0.142857143\\ 0.214285714 & 0.642857143 & 0 & -0.428571429\\ 0 & 0 & 0 & 0\\ -0.142857143 & -0.428571429 & 0 & 0.285714286\end{pmatrix}\begin{pmatrix}1\\ 0\\ 0\\ 0\end{pmatrix}$$

$$= \begin{pmatrix}0.06287143\\ 0.208571429\\ -0.01\\ -0.155714286\end{pmatrix}.$$

For

$$w = \begin{pmatrix}10\\ 10\\ 10\\ 10\end{pmatrix}$$

we obtain

$$\begin{pmatrix}\beta_{41}\\ \beta_{42}\\ \beta_{43}\\ \beta_{44}\end{pmatrix} = \begin{pmatrix}1.42\\ 4.28\\ -0.01\\ -2.87\end{pmatrix}.$$

All these solutions, for arbitrary vectors $w$, are consistent with the first four survival vectors. The space they span is only guaranteed to include the fifth vector if $w$ happens to be correct, but the values of its components are *a priori* unknown.

It is not clear *per se* how to choose this vector $w$. One criterion to be enforced could be that $\beta_{ii}$ should be negative, because it is related to the carrying capacity and responsible for slowing down the growth of population 1. In fact, for a single population, the LV system reduces to the well-known logistic growth law, where this parameter is always negative.

### 3. Further Details of the Analysis of a Bacterial Community with Four Species [1]

#### *3.1. Biological details associated with the bacterial community*

The data for the ALVI analysis in the Text come from a recent study of four bacterial species grown in Castrol Metal-Working Fluid (MWF) [1]. The species (with abbreviations we use) were *Agrobacterium tumefaciens* (*At*), *Comamonas testosteroni* (*Ct*), *Microbacterium saperdae* (*Ms*), and *Ochrobactrum anthropi* (*Oa*). The study was designed to characterize how toxicity may drive the interactions between bacterial species. Specifically, the authors examined growth in three media: MWF with and without supplemental casamino acids, consisting of a mixture of amino acids with very small peptides obtained from acid hydrolysis of casein, and an amino acid medium without MWF. All species were first grown by themselves in monoculture, and then together with one or more other species as pairs or in combinations of three. Finally, the community of all four species was analyzed. All communities were studied in triplicate in 30mL of medium, and a 200μL aliquot was taken each day for 6 days and then on days 8 and 12 to quantify the abundances of all species in each given medium. The counting was accomplished with selective plating, that is, in specific media or with antibiotics that permitted just one of the species to survive. The results consisted of densities, expressed as colony-forming units per milliliter (CFU/mL). The authors calculated interspecies interactions by first calculating the area under the curve (AUC) of each species' growth on its own, and then comparing it to the AUC of the growth in the presence of another species. The authors found that the species were only cooperative with each other when grown in the MWF. However, when supplemental amino acids were added or MWF removed, the species became competitive with one another, showing that the environment strongly contributes to interspecies interactions.

Here we use the community of all four species, grown on the amino acid medium without MWF, as shown in Figure S8 of the original paper. As this figure indicates, two of the populations begin to decline about 150 hours into the experiments, presumably due to the depletion of nutrients.

### *3.2. Splines*

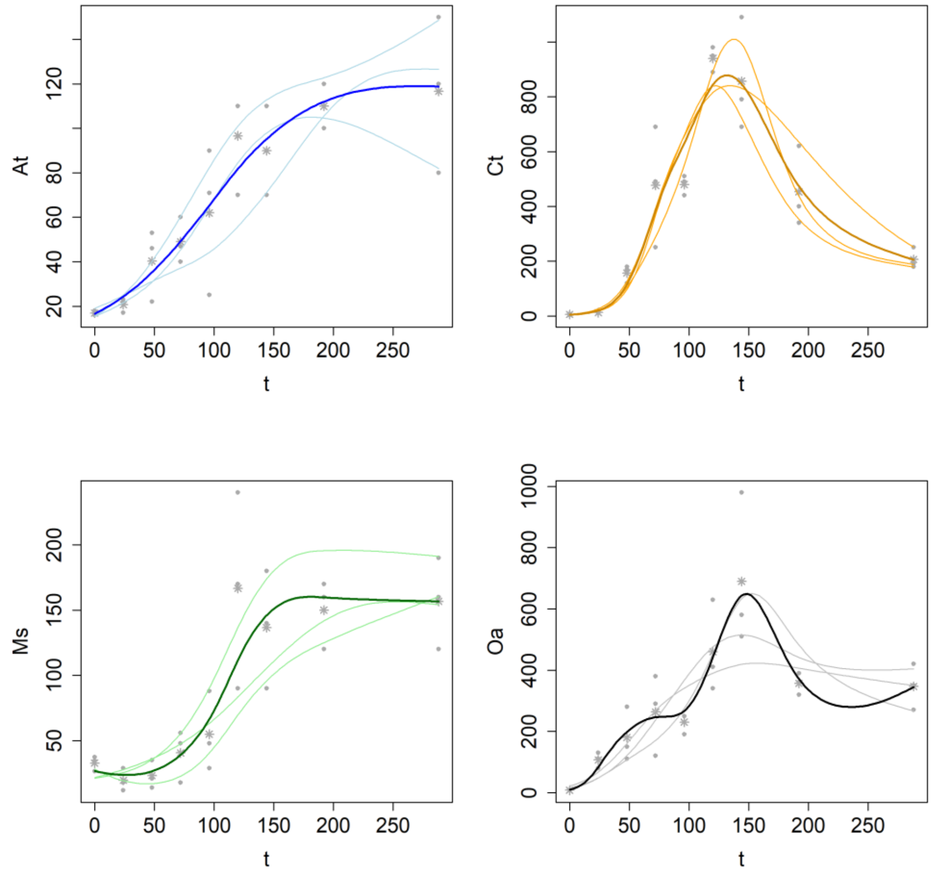


**Fig. S4: Smoothing splines of trends of four co-cultured bacterial populations [1], in units of 1,000,000.** Each trend was measured in three replicates. Data ●: replicates; light curves; ✳: means of three replicates; light curves: splines of individual replicates; dark curves: splines of replicate means. Species abbreviations: *Agrobacterium tumefaciens* (*At*), *Comamonas testosteroni* (*Ct*), *Microbacterium saperdae* (*Ms*), and *Ochrobactrum anthropi* (*Oa*).

### *3.3. Details of ALVI Application*

**Table S2: Degrees of freedom and point samples for fitting the dynamics of a community of four bacterial species described in [1] with LV models.**

| | ***At*** | ***Ct*** | ***Ms*** | ***Oa*** | **Point Sample** |
|---|---|---|---|---|---|
| **Replicate 1** | 4 | 5 | 3 | 4 | *t* = 24, 72, 96, 144, 288 |
| **Replicate 2** | 4 | 6 | 4 | 4 | *t* = 24, 72, 120, 192, 288 |
| **Replicate 3** | 4 | 6 | 5 | 5 | *t* = 24, 72, 96, 144, 192 |
| **Means** | 4 | 6 | 5 | 7 | *t* = 24, 72, 120, 144, 288 |

### Table S3: Parameter Values of Fits to Individual Replicates and their Mean

| | Replicate 1 | Replicate 2 | Replicate 3 | Mean of Replicates |
|---|---|---|---|---|
| $a_1$ | 0.0142402 | 0.0224366 | 0.0162296 | 0.0196418 |
| $b_{11}$ | -0.0004535 | 0.0000251 | 0.0001256 | -0.0001369 |
| $b_{12}$ | 0.0000093 | 0.0000068 | -0.0000055 | 0.0000026 |
| $b_{13}$ | 0.0003619 | -0.0000585 | -0.0002532 | -0.0000254 |
| $b_{14}$ | -0.0000432 | -0.0000343 | 0.0000084 | -0.0000001 |
| $a_2$ | 0.0871680 | 0.0668200 | 0.1095992 | 0.0920004 |
| $b_{21}$ | -0.0036469 | 0.0002872 | -0.0024362 | -0.0011569 |
| $b_{22}$ | -0.0001442 | 0.0000304 | 0.0001718 | -0.0000114 |
| $b_{23}$ | 0.0023897 | -0.0002292 | 0.0016731 | 0.0003807 |
| $b_{24}$ | 0.0000909 | -0.0001852 | -0.0003276 | -0.0000524 |
| $a_3$ | 0.0129389 | 0.0135113 | -0.0279418 | -0.0063944 |
| $b_{31}$ | -0.0000061 | -0.0000655 | 0.0018354 | 0.0007739 |
| $b_{32}$ | 0.0000048 | -0.0000386 | -0.0000407 | 0.0000324 |
| $b_{33}$ | -0.0001116 | -0.0004112 | -0.0012545 | -0.0005752 |
| $b_{34}$ | 0.0000093 | 0.0002009 | 0.0000287 | -0.0000067 |
| $a_4$ | 0.0562003 | 0.0468287 | 0.0668262 | 0.1057096 |
| $b_{41}$ | -0.0013472 | 0.0002627 | -0.0016605 | -0.0027523 |
| $b_{42}$ | -0.0000205 | 0.0000840 | 0.0001588 | 0.0000397 |
| $b_{43}$ | 0.0010468 | 0.0002128 | 0.0013359 | 0.0019270 |
| $b_{44}$ | -0.0001243 | -0.0003488 | -0.0002897 | -0.0002389 |

**Table S4: LV Parameter Values of Fits for Replicate Means Spanning Different Time Intervals. Plots of Corresponding Trajectories are shown in Figures 3 and 4 of the Text**

| | $t \in [0, 288]$ | $t \in [0, 144]$ | Absolute Difference |
|---|---|---|---|
| $a_1$ | 0.019642 | 0.011071 | 0.00857 |
| $b_{11}$ | -0.00014 | 0.000373 | -0.00051 |
| $b_{12}$ | 2.6E-06 | -2.1E-05 | 2.37E-05 |
| $b_{13}$ | -2.5E-05 | -7.6E-05 | 5.01E-05 |
| $b_{14}$ | -1E-07 | -1.9E-05 | 1.85E-05 |
| $a_2$ | 0.092 | -0.1067 | 0.198696 |
| $b_{21}$ | -0.00116 | 0.009878 | -0.01103 |
| $b_{22}$ | -1.1E-05 | -0.00053 | 0.000522 |
| $b_{23}$ | 0.000381 | -0.00071 | 0.001095 |
| $b_{24}$ | -5.2E-05 | -0.0004 | 0.000344 |
| $a_3$ | -0.00639 | -0.01422 | 0.007821 |
| $b_{31}$ | 0.000774 | 0.001037 | -0.00026 |
| $b_{32}$ | 3.24E-05 | 2.6E-06 | 2.98E-05 |
| $b_{33}$ | -0.00058 | -0.00051 | -6.6E-05 |
| $b_{34}$ | -6.7E-06 | 5.2E-06 | -1.2E-05 |
| $a_4$ | 0.10571 | 0.078073 | 0.027636 |
| $b_{41}$ | -0.00275 | -0.00175 | -0.001 |
| $b_{42}$ | 3.97E-05 | 7.3E-06 | 3.24E-05 |
| $b_{43}$ | 0.001927 | 0.001076 | 0.000851 |
| $b_{44}$ | -0.00024 | -0.0001 | -0.00014 |

**Reference**

[1] Piccardi, P., B. Vessman, and S. Mitri, Toxicity drives facilitation between 4 bacterial species. P*NAS U.S.A.*, **116**(32), 15979-15984, 2019.
